# Reciprocal regulation between the protein arginine deiminases and mSWI/SNF chromatin remodelers controls skeletal muscle differentiation and regeneration

**DOI:** 10.64898/2026.01.20.700682

**Authors:** Monserrat Olea-Flores, Tapan Sharma, Anand Parikh, Leonard Barasa, Sarah Hachmer, F. Jeffrey Dilworth, Teresita Padilla-Benavides, Paul R. Thompson, Anthony N. Imbalzano

## Abstract

Protein arginine deiminases (PADs) post-translationally convert arginine to citrulline on target proteins and serve as regulators of multiple cellular functions. PAD enzymes have been implicated in autoimmune disorders, cancer, and other diseases. Inhibiting PAD activity is currently being pursued clinically for therapeutic purposes. However, little is known about PAD function in normal developmental or homeostatic processes. Here we show that multiple PAD isoforms contribute to primary myoblast differentiation by binding to regulatory regions of target genes. Furthermore, we demonstrate a novel, reciprocal requirement for PADs and mammalian SWI/SNF (mSWI/SNF) chromatin remodeling enzymes; PAD enzymes are required for the expression and binding of specific mSWI/SNF enzyme subunits to target gene regulatory sequences while mSWI/SNF enzymes are required for the expression and binding of PAD enzymes. *In vivo*, the PADs contribute to mouse skeletal muscle regeneration after injury, with PAD4 specifically identified as a required regulator. This work identifies the PADs as critical cofactors in the initiation of skeletal muscle differentiation and reveals previously unappreciated connections between two major co-activator families during normal tissue development. Moreover, the results reveal important considerations for ongoing therapeutic approaches to myriad human diseases that utilize inhibitors of each enzyme family.

## INTRODUCTION

Studies of skeletal muscle development during embryogenesis and in adult muscle growth and regeneration have revealed basic principles of cell differentiation, cell cycle control, and gene regulation. For example, molecular dissection of skeletal muscle differentiation has demonstrated the complex interplay between transcription factors, co-regulators, including chromatin modifiers and remodelers, and signaling molecules^1–5^. These factors control the ability of a precursor cell to initiate and execute a new program of gene expression to create differentiated tissue. Prior work has shown how mSWI/SNF chromatin remodeling enzymes, histone acetyl transferases (HAT) and deacetylases (HDAC), lysine and arginine methyltransferases, kinases, and other cofactors that impact chromatin structure and function affect myogenic transcription factor function at enhancers and promoters of genes that are differentially expressed during myoblast differentiation. Mechanistic studies have demonstrated the functions of different factors and chromatin alterations and modifications at target genes^6–15^, while also exploring the impact of three-dimensional organization of the genome during differentiation^16–20^.

Protein arginine deiminases (PADs) are enzymes that convert arginine into citrulline on protein targets found throughout the cell^21,22^. Histones are among the substrates, and citrullination occurs on linker histone and at about two dozen sites in the core histones^23,24^. The H3 and H4 tail residues H3-R2,-R8,-R17,-R26, and H4-R3 have been best studied^23,25^. These post-translational histone modifications are generally associated with gene activation, as removal of the positive charge on arginine is thought to promote chromatin decompaction^23^. There is little information about how PADs affect cellular differentiation. How PAD enzymes integrate with other regulatory cofactors is also poorly understood.

Here, we identify the PAD family of transcriptional co-regulators as a previously unappreciated contributor to myoblast differentiation in culture and *in vivo*. We also demonstrate a previously unidentified regulatory dependency between the PADs and ATP-dependent chromatin remodeling enzymes of the mammalian SWI/SNF (mSWI/SNF) family. mSWI/SNF enzyme subunits have been reported to be mutated and/or mis-regulated in 20% of all human cancers^26^ and play a role in neurodevelopmental diseases^27–29^. PAD enzyme mis-regulation is associated with autoimmune diseases and in cancers^21,22,30,31^. The newly reported relationship between mSWI/SNF and PAD enzymes suggests that therapeutic approaches for diseases caused by one enzyme family may be relevant for diseases caused by either.

## RESULTS

### Multiple PAD enzymes promote myoblast differentiation and myogenic gene expression

Even though a PAD enzyme was originally purified from skeletal muscle^32^, little is known about the role of the PADs in the normal formation or function of skeletal muscle is known. We initially assessed PAD gene expression in an RNA-seq dataset comparing proliferating and differentiating immortalized C2C12 myoblasts^33^. The analysis showed that PAD2, PAD3, and PAD4 gene expression was elevated in myoblasts that were differentiated compared to expression in proliferating myoblasts (**Supp. Fig. 1A**).

Validation by RT-qPCR and immunoblotting confirmed increased PAD expression during differentiation (**Supp. Fig. 1B-D**) as well as a corresponding increase in citrullinated H3R17 (**Supp. Fig. 1E**). Importantly, we demonstrated that this differentiation-dependent induction of PAD2, PAD3, and PAD4 was recapitulated in primary myoblasts derived from murine satellite cells, which are adult stem cells for skeletal muscle^34^, at both the mRNA and protein levels (**Fig. 1**).

**Figure 1.**
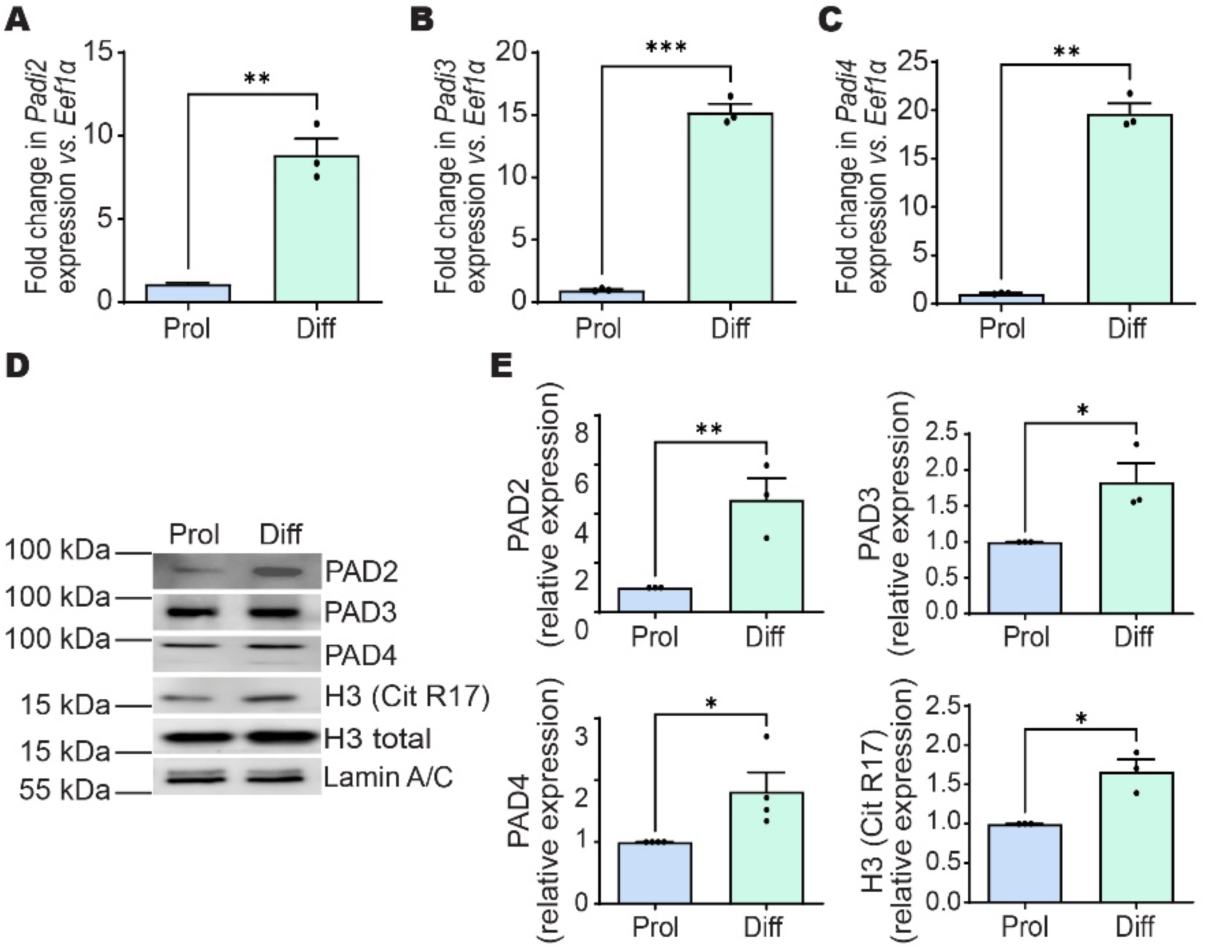
**Induction of PAD2, PAD3, and PAD4 expression during myoblast differentiation**. **(A-C)** Relative mRNA levels of *Padi2* **(A)**, *Padi3* **(B)**, and *Padi4* **(C)** were measured by qPCR in primary myoblasts under proliferating conditions (Prol) or after 24 h of differentiation (Diff). **(D)** Representative immunoblots showing PAD2, PAD3, PAD4, and citrullinated histone H3 (H3 CitR17) protein levels in proliferating and differentiating primary myoblasts. **(E)** Quantification of PAD2, PAD3, PAD4 protein levels normalized to Lamin A/C levels and H3 CitR17 protein abundance normalized to total histone H3. Data are presented as mean ± SEM from three independent experiments with *p < 0.05, **p < 0.01, and ***p < 0.001.

To directly assess the requirement for these PADs, we knocked down each of these isoforms by stably expressing one of two different shRNAs in murine primary myoblasts (**Fig 2A-C**). Western blot (left) and quantitative PCR analysis (right) showed that the knockdowns were specific, with each shRNA affecting only the targeted PAD protein and mRNA. Immunoblot analyses also demonstrated that each PAD knockdown reduced the expression of myogenin, a major myogenic transcription factor that drives differentiation^35–38^, and the differentiation marker myosin heavy chain (MHC; **Fig. 2A-C**), suggesting a differentiation defect. Comparison of the differentiation capacity of wild-type (WT) and scrambled (Scr) shRNA control myoblasts with *Padi2*, *Padi3*, or *Padi4* knockdown myoblasts confirmed that knockdown of each PAD impaired differentiation and reduced expression of both differentiation markers (**Fig. 2D,E)**. Consistent with results in primary myoblasts, knockdown of *Padi2*, *Padi3*, or *Padi4* in immortalized C2C12 myoblasts impaired differentiation (**Supp. Fig. 1F-I**). Together, the data indicate that each of the three PAD enzymes contributes to myoblast differentiation.

**Figure 2.**
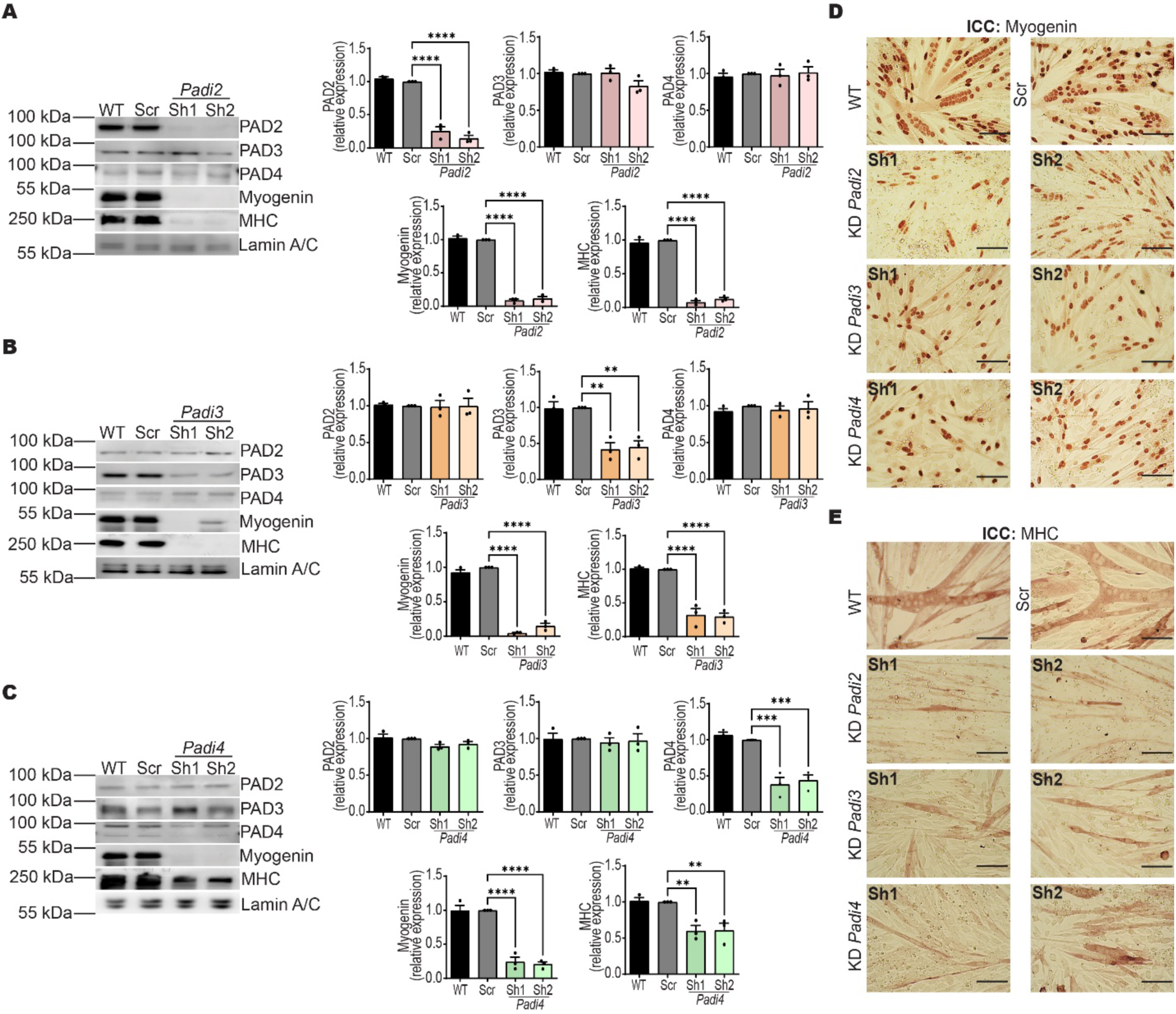
Knockdown of PAD2, PAD3, or PAD4 compromises myoblast differentiation. (A-C) Representative immunoblots (left) and corresponding quantification (right) of PAD2, PAD3, PAD4, myogenin, and myosin heavy chain (MHC) protein levels in primary myoblasts differentiated for 24 h following shRNA-mediated knockdown of *Padi2* **(A)**, *Padi3* **(B)**, or *Padi4* **(C)** using two different shRNA sequences (Sh1 and Sh2). Lamin A/C was used as a loading control. Bar graphs represent mean ± SEM from three independent experiments; **p < 0.01, ***p < 0.001, and ****p < 0.0001. Scr, scrambled sequence shRNA; WT, wild type. **(D,E)** Representative phase-contrast micrographs of primary myoblasts differentiated for 24 h (WT, Scr, *Padi2, Padi3*, or *Padi4* knockdown (KD)) and immunostained for myogenin **(D)** or MHC **(E)**. Scale bar, 100 μm.

As suggested by the western blot and immunohistochemistry analyses, the genes encoding *Myogenin* and myosin heavy chain IIB (*MyhIIb*), a myosin heavy chain isoform expressed in fast-twitch muscles of rodents^39^, were downregulated as a consequence of *Padi* gene knockdown (**Supp. Fig. 2**). However, assessment of additional individual genes impacted by each *Padi* gene knockdown showed that not all genes associated with myogenesis were affected equivalently. For instance, the expression of the myomaker (*Mymk*) and muscle creatine kinase (*Ckm*) genes were affected by *Padi2* and *Padi4* knockdown but not by *Padi3* knockdown, whereas caveolin 3 (*Cav3*) expression was affected by *Padi2* and *Padi3* knockdown, but not by *Padi4* knockdown (**Supp. Fig. 2)**.

### PAD2, PAD3, and PAD4 exhibit dual cytoplasmic-nuclear localization consistent with a role in transcriptional regulation

To assess the potential mechanisms by which PAD2, PAD3, and PAD4 regulate myogenic gene expression and differentiation, we first determined their subcellular localization. Previous studies indicated that while PAD2 is predominantly cytoplasmic, it can translocate into the nucleus in response to Ca^2+^ signaling^40^. There, it is known to citrullinate histones, such as H3R26, to regulate gene expression^40^. PAD4 localizes to both the cytoplasm and nucleus as it has a nuclear localization signal that is essential for H2, H3 and H4 citrullination^24,30,41–45^. PAD3 has only been reported to be a cytosolic enzyme^46^.

Using complementary immunofluorescence (**Fig. 3A**) and biochemical fractionation approaches (**Fig. 3B**), we found that these three PAD enzymes are present in both the cytoplasmic and nuclear compartments. This dual localization indicates that PAD2, PAD3, and PAD4 have the potential to directly access chromatin and participate in transcriptional regulation during myoblast differentiation.

**Figure 3.**
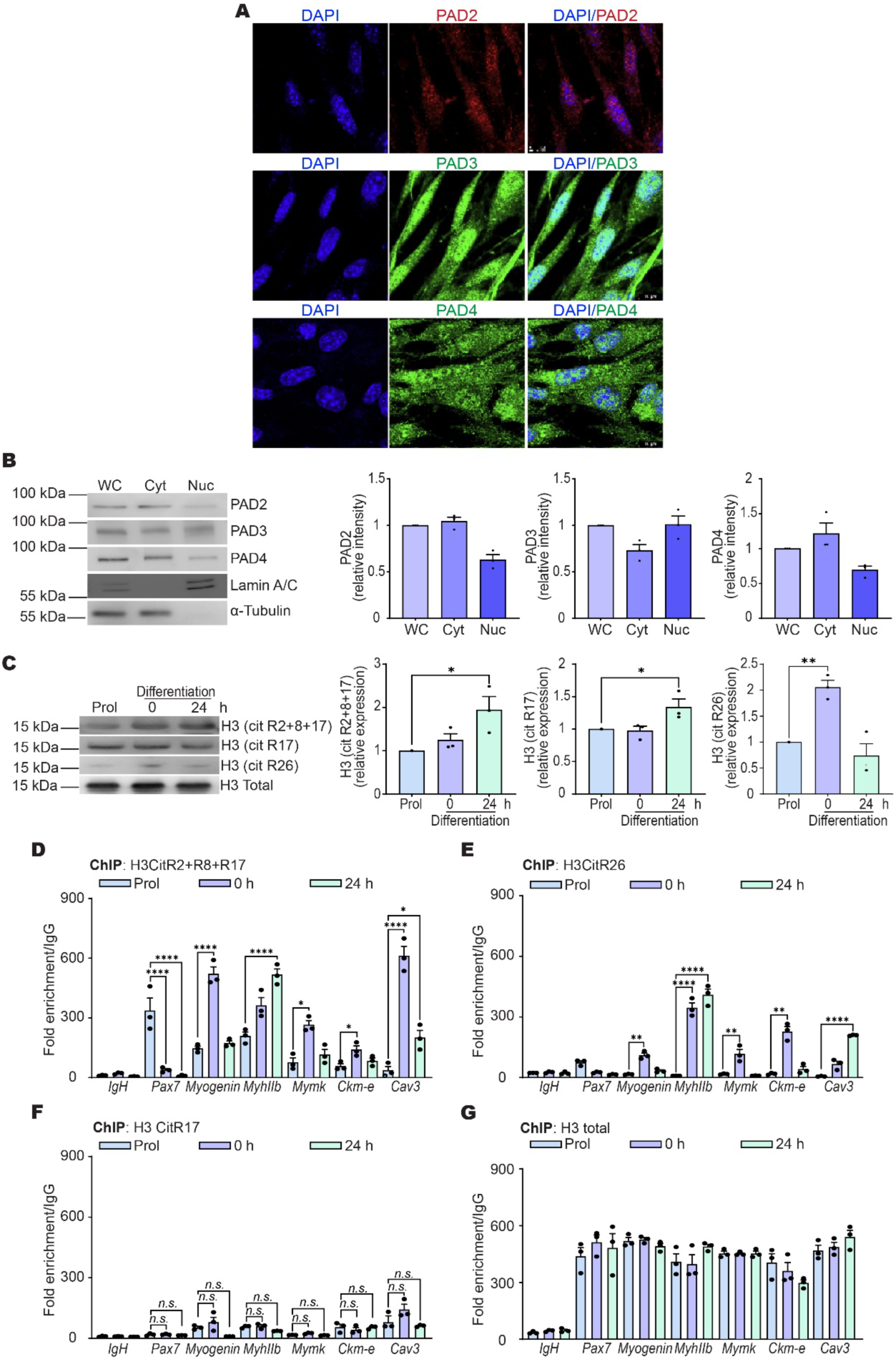
PAD2, PAD3, and PAD4 localize to the cytoplasm and the nucleus in differentiating myoblasts. Histone citrullination increases both in bulk and at myogenic gene regulatory sequences during differentiation. **(A)** Confocal micrographs of primary myoblasts differentiated for 24 h and immunostained with anti-PAD2 (red) and anti-PAD3 or anti-PAD4 (green) antibodies. Nuclei were counterstained with DAPI (blue). Scale bar, 5 μm. **(B)** Subcellular fractionation of primary myoblasts differentiated for 24 h showing whole-cell (WC), cytosolic (Cyt), and nuclear (Nuc.) fractions. Representative immunoblots (left) and corresponding quantification (right) of PAD2, PAD3, and PAD4 protein levels are shown. Lamin A/C and α-tubulin were used as purity controls for the nuclear and cytoplasmic fractions, respectively. 20 ug of each extract was loaded onto the gels. **(C)** Representative immunoblots (left) and quantification (right) of histone H3 citrullination marks (H3CitR2+R8+R17, H3CitR17, and H3CitR26) in proliferating (P or Prol), time 0 (0 h), or differentiating (24 h) primary myoblasts. **(D-G)** ChIP-qPCR analysis showing enrichment of H3CitR2+R8+R17 **(D)**, H3CitR26 **(E)**, H3CitR17 **(G)** and total H3 **(H)** at the *Pax7* promoter and at regulatory sequences controlling the indicated myogenic genes in proliferating (Prol), time 0 (0 h), and differentiating (24 h) primary myoblasts. The IgH enhancer was included as a negative sequence control. Bar graphs represent mean ± SEM from three independent experiments. *p < 0.05, **p < 0.01, ***p < 0.001, and ****p < 0.0001. n.s., not significant.

### Histone citrullination increases during myoblast differentiation and correlates with PAD expression

PADs citrullinate a wide range of protein substrates^21,22^, including nuclear targets such as linker and core histones^41,47,48^. Histone citrullination has generally been associated with increased chromatin accessibility and transcriptional competence^49^. Given the nuclear localization of PAD2, PAD3, and PAD4, we next examined whether histone citrullination changes during myoblast differentiation. To this end, we assessed levels of citrullinated H3 in proliferating myoblasts, at the onset (0 h) and at the completion of differentiation (24 h). Immunoblotting with an antibody recognizing H3 citrullinated at R2, R8, and R17 revealed no significant change between proliferating myoblasts and those at differentiation onset, but an approximately two-fold increase in differentiated cells (**Fig. 3C**). Analysis of H3 cit-R17 alone showed a similar, though more modest, increase at 24 h (**Fig. 3C**). By contrast, H3 cit-R26 exhibited a distinct temporal pattern, with a marked increase at the onset of differentiation followed by a return to baseline levels in differentiated myotubes (**Fig. 3C**). Notably, the sustained increase in H3 cit-R2/8/17 and H3 cit-R17 coincided with the differentiation-associated upregulation of PAD2, PAD3, and PAD4 protein levels, whereas the transient induction of H3 cit-R26 may suggest a potential role for this modification during earlier stages of lineage commitment.

To determine whether these citrullinated histones are incorporated into myogenic gene regulatory regions, we performed chromatin immunoprecipitation analyses at multiple differentiation-associated gene regulatory sequences (**Fig. 3D-G**). Regulatory regions of the examined genes that are activated during differentiation showed increased incorporation of H3 cit-R2/8/17 (**Fig. 3D**) and H3 cit-R26 (**Fig. 3E**) during differentiation, although the timing and magnitude of enrichment varied between genes. By contrast, incorporation of H3 cit-R17 (**Fig. 3F**) alone did not change significantly over the course of differentiation, despite its increased global abundance. Total H3 was used as the control (**Fig. 3G**). Notably, the incorporation of H3 cit-R2/8/17 and H3 cit-R26 decreased at the *Pax7* promoter as a function of differentiation (**Fig. 3D,E**). *Pax7* is a marker of proliferating myoblasts; expression decreases as differentiation proceeds^50^.

These data indicate that multiple arginine residues within the H3 tail undergo citrullination during myoblast differentiation, that global increases in histone citrullination correlate with rising PAD expression, and that citrullinated histones are selectively incorporated into myogenic gene regulatory regions in a manner consistent with transcriptional activity.

We then examined incorporation of H3 cit-R2/8/17 and H3 cit-R26 in primary myoblasts in which *Padi2*, *Padi3*, or *Padi4* was knocked down (**Supp. Fig. 3A-C**). Consistent with the limited understanding of PAD isoform specificity toward individual histone residues, the effects of PAD depletion were both gene-and modification-specific. No single PAD enzyme could be definitively linked to the incorporation of a specific citrullinated H3 residue at myogenic loci. These findings suggest functional redundancy and/or coordinated activity among PAD family members in shaping the histone citrullination landscape during myoblast differentiation, rather than a strict one enzyme-one modified residue model. However, despite the possibility of redundancy in modifying specific histone, the data clearly show that the PAD enzymes are not redundant in promoting myoblast differentiation.

### PADs 2, 3, and 4 occupy regulatory chromatin in a transcription-dependent manner

One of the most direct mechanisms by which a PAD could regulate myogenic gene expression is through physical association with chromatin at transcriptionally active or poised loci. To test this possibility, we examined PAD2, PAD3, and PAD4 binding at myogenic gene regulatory sequences. All three PAD enzymes were enriched at the regulatory sequences of the genes examined for histone citrullination, except for the *Mymk* promoter, which was not occupied by PAD3 (**Fig. 4A-C**). PAD2, PAD3, and PAD4 were robustly associated with the *Pax7* promoter in proliferating myoblasts, but this binding was lost as differentiation progressed (**Fig. 4A-C**). Together, these results demonstrate that PAD2, PAD3, and PAD4 interact directly with chromatin at regulatory regions of differentiation-dependent genes, and that PAD binding closely tracks transcriptional activity, supporting a role for the PADs as dynamic chromatin-associated regulators of gene expression during myoblast differentiation.

**Figure 4.**
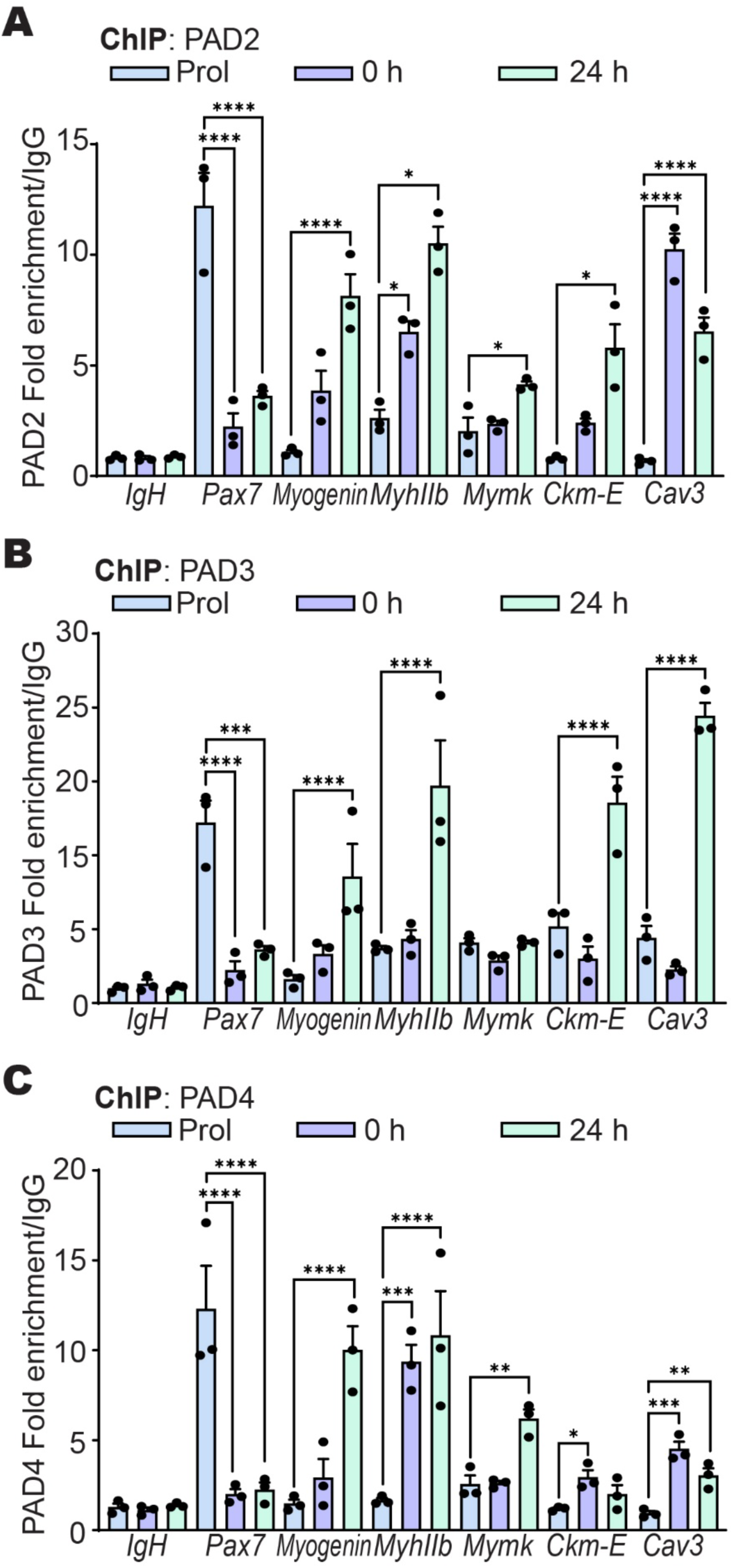
PAD2, PAD3, and PAD4 occupy myogenic gene regulatory elements during differentiation. ChIP-qPCR analysis showing enrichment of PAD2 **(A)**, PAD3 **(B)**, and PAD4 **(C)** at the *Pax7* promoter and at regulatory sequences controlling the indicated myogenic genes in primary myoblasts under proliferating conditions (Prol), at time 0 (0 h), and after 24 h of differentiation. The IgH enhancer was included as a negative sequence control. Bar graphs represent the mean ± SEM from three independent experiments. *p < 0.05, **p < 0.01, ***p < 0.001, and ****p < 0.0001.

### PAD enzymes enable the recruitment of the cBAF subfamily of mSWI/SNF chromatin remodeling enzymes to myogenic gene regulatory sequences by regulating chromatin remodeler subunit expression

A defining feature of myogenic gene activation during myoblast differentiation is the recruitment of ATP-dependent mSWI/SNF chromatin-remodeling complexes to myogenic regulatory elements^6,7,12^, particularly the canonical BAF (cBAF) subfamily^51^. The cBAF-specific subunits BAF250A (ARID1A) and DPF2 distinguish this complex from other mSWI/SNF assemblies and are essential for myogenic transcriptional activation^52^. Knockdown of any *Padi* gene, in general, reduced BAF250A and DPF2 occupancy at myogenic gene regulatory sequences (**Fig. 5A,B**). A few gene-and PAD isoform-exceptions existed. For example, *Padi4* knockdown did not significantly diminish BAF250A binding at the *Myogenin* promoter (**Fig. 5A**), and neither *Padi2* nor *Padi3* knockdown significantly reduced DPF2 binding at the *Mymk* promoter (**Fig. 5B**). The reasons for the PAD-and locus-specific differences are not understood, though we can propose localized redundancy by individual PAD enzymes.

**Figure 5.**
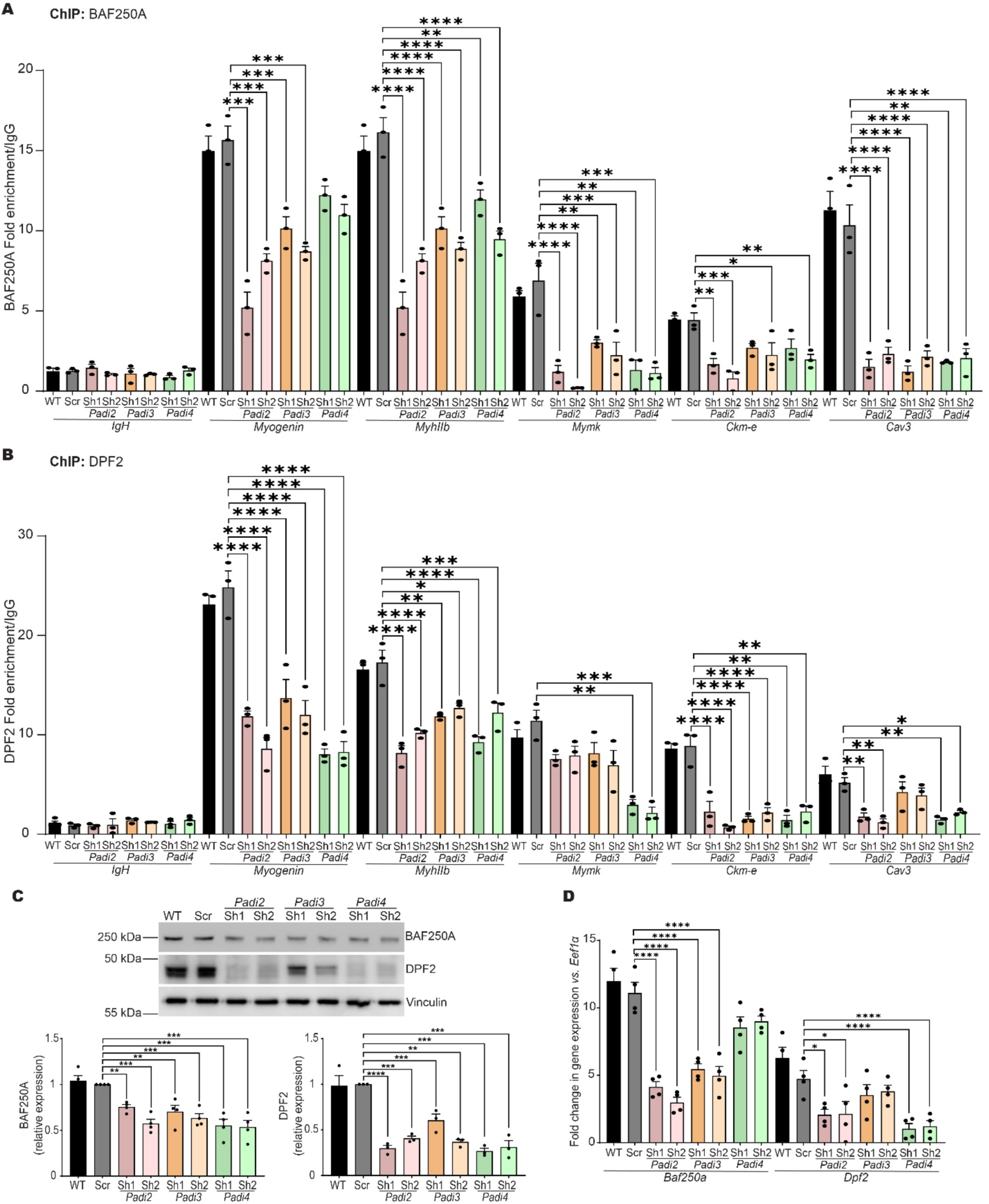
Binding of the cBAF-specific mSWI/SNF chromatin remodeling enzyme subunits BAF250A and DPF2 to myogenic gene regulatory sequences is compromised by *Padi* gene knockdown. *Padi* gene knockdown reduces the levels of BAF250A and DPF2. (A,B) ChIP-qPCR analysis showing enrichment of the cBAF subunits BAF250A **(A)** and DPF2 **(B)** at regulatory sequences of the indicated myogenic genes in primary myoblasts differentiated for 24 h following shRNA-mediated knockdown of *Padi2, Padi3*, or *Padi4*. Distinct shRNAs for each *Padi* gene are indicated by Sh1 and Sh2. **(C)** Representative immunoblots (top) and corresponding quantification (bottom) of BAF250A and DPF2 protein levels in primary myoblasts differentiated for 24 h under *Padi2, Padi3*, or *Padi4* knockdown conditions. Vinculin was used as a loading control. **(D)** Relative mRNA levels of *Baf250a* and *Dpf2* measured by qPCR in primary myoblasts following 24 h of differentiation. The IgH enhancer was included as a negative sequence control. Bar graphs represent the mean ± SEM from three independent experiments. *p < 0.05, **p < 0.01, ***p < 0.001, and ****p < 0.0001. WT, Wild-type; Scr, scrambled sequence shRNA.

The data indicate that PAD2, PAD3, and PAD4 each can contribute to efficient recruitment of cBAF subunits to myogenic regulatory sequences. To investigate the basis for impaired cBAF binding, we examined cBAF subunit expression following *Padi* gene knockdown. Immunoblot analysis revealed that depletion of *Padi2*, *Padi3*, or *Padi4* reduced BAF250A and DPF2 protein levels (**Fig. 5C**). However, *Baf250A* mRNA levels were reduced only upon *Padi2*-or *Padi3*-knockdown, while *Dpf2* mRNA levels were only significantly reduced upon knockdown of *Padi2* or *Padi4* (**Fig. 5D**), suggesting contributions from both transcriptional and post-transcriptional regulatory mechanisms. Collectively, these results demonstrate that PAD enzymes are required for efficient cBAF occupancy at myogenic regulatory sequences, at least in part by maintaining appropriate expression levels of key chromatin-remodeling subunits, thereby enabling productive myogenic gene activation.

### Reciprocal regulation between PAD enzymes and cBAF mSWI/SNF complexes coordinates their binding to myogenic chromatin

Our previous results suggested that PAD enzymatic activity might function upstream of mSWI/SNF chromatin remodeling during myogenic gene activation. If this were the case, depletion of mSWI/SNF subunits would be predicted to have little or no effect on PAD2, PAD3, or PAD4 binding to myogenic gene regulatory regions. However, experimental testing of this hypothesis did not support this model. Knockdown of the cBAF subunits *Baf250a* or *Dpf2* inhibited myoblast differentiation, consistent with our prior published results in immortalized C2C12 myoblasts^51^ (**Supp. Fig. 4A**). Knockdown of *Baf250a* (**Fig. 6A-C**) or *Dpf2* (**Fig. 6D-F**) resulted in a reduction in PAD2 and PAD4 occupancy at both the *myogenin* and *MyhIIb* promoters. Knockdown of *Baf250a* reduced PAD3 binding at the *myogenin* promoter but not the *MyhIIb* promoter (**Fig. 6B**), while knockdown of *Dpf2* reduced PAD3 binding at the *MyhIIb* promoter but not the *myogenin* promoter (**Fig. 6E**). PAD2 occupancy at other myogenic regions (*Mymk*, the *Ckm* enhancer, and *Cav3*) was significantly reduced in both *Baf250A*-and *Dpf2*-knockdown cells (**Supp. Fig. 4B,C**). PAD3 and PAD4 binding at these loci showed gene-specific results (**Supp. Fig. 4D-G**). The molecular basis for gene-and cBAF subunit-specific exceptions remains unknown, but it is possible that at some loci, knockdown of only one subunit of the cBAF complex does not impact function enough to register in this binding assay. Nonetheless, these data demonstrate that PAD enzyme association with myogenic regulatory sequences is frequently compromised in the absence of normal levels of BAF250A or DPF2, indicating that mSWI/SNF complexes are generally required for PAD chromatin binding. Furthermore, these data provide additional mechanistic support for our prior results indicating the cBAF complex is essential for myoblast differentiation^51^.

**Figure 6.**
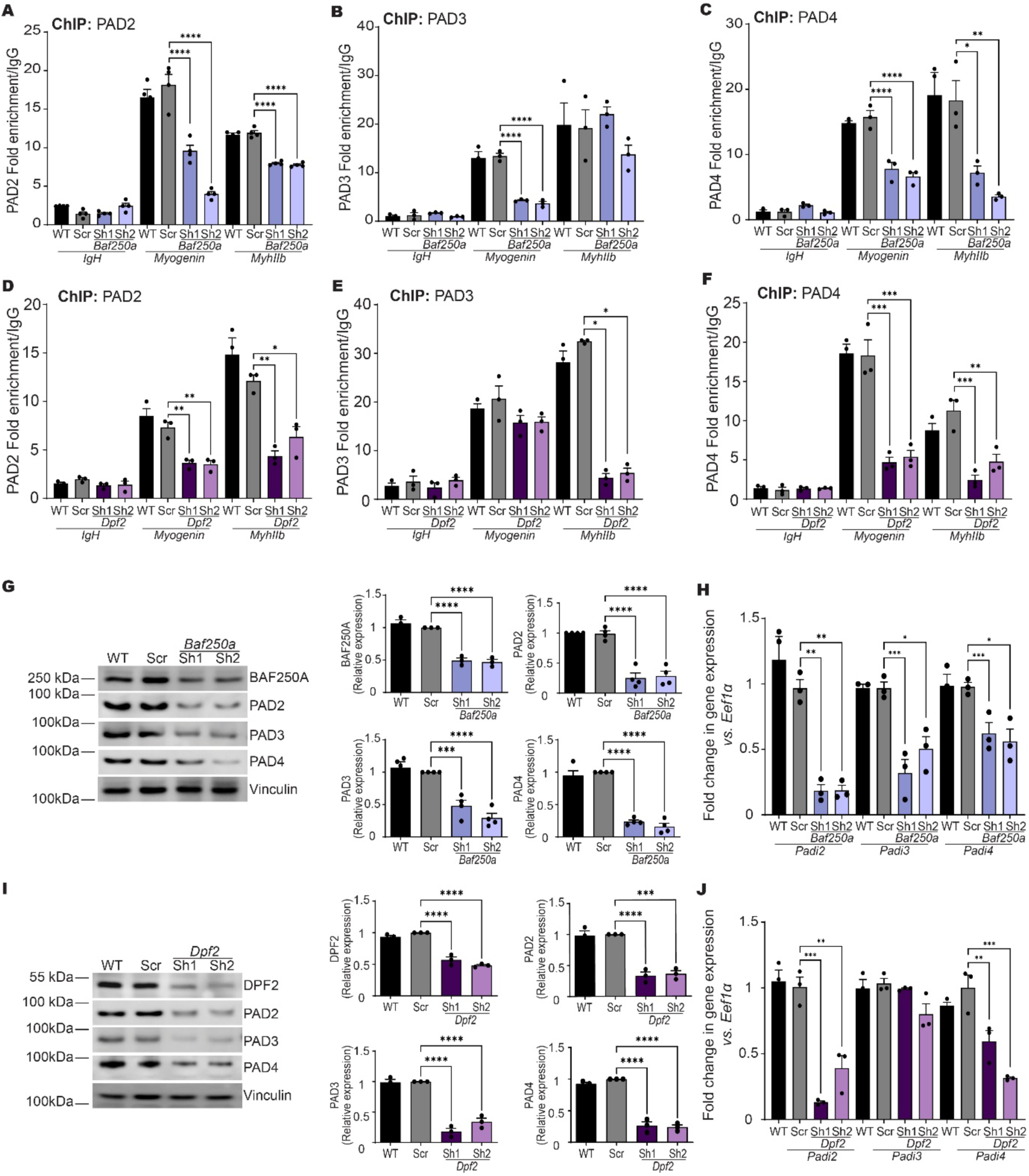
Knockdown of the genes encoding cBAF-specific mSWI/SNF chromatin remodeling enzyme subunits BAF250A and DPF2 reduces the binding of PAD2, PAD3, and PAD4 to myogenic gene regulatory sequences in differentiating myoblasts. Knockdown of *Baf250A* or *Dpf2* reduces the levels of the PADs. (A-C) ChIP-qPCR analysis showing enrichment of PAD2, PAD3, and PAD4, respectively, at the *IgH* (negative control), *Myogenin*, and *MyhIIb* promoters in differentiating primary myoblasts following *Baf250a* knockdown. Distinct shRNAs for *Baf250A* are indicated by Sh1 and Sh2 **(D-F)** As in (A-C) except following *Dpf2* knockdown. **(G, I)** Representative immunoblots (left) and quantification (right) of PAD2, PAD3, and PAD4 protein levels in primary myoblasts differentiated for 24 h following *Baf250a* knockdown or *Dpf2* knockdown, respectively. Vinculin was used as a loading control. **(H, J)** Relative mRNA levels of *Padi2, Padi3*, and *Padi4* measured by qPCR in primary myoblasts differentiated for 24 h following knockdown of *Baf250a* or *Dpf2*, respectively. Bar graphs represent mean ± SEM from three independent experiments; *p < 0.05, **p < 0.01, ***p < 0.001, and ****p < 0.0001. Scr, scrambled sequence shRNA; WT, wild type.

Immunoblot analysis confirmed that knockdown of *Baf250A* and *Dpf2* in the respective cell populations (**Fig. 6G,I**) also reduced the levels of PAD2, PAD3, and PAD4 proteins. This suggests that diminished PAD chromatin binding was at least partially due to decreased PAD protein abundance. Consistent with this interpretation, analysis of transcript levels revealed that *Baf250a* knockdown significantly reduced expression of *Padi2*, *Padi3*, and *Padi4* mRNAs (**Fig. 6H**). *Dpf2* knockdown selectively reduced *Padi2* and *Padi4* mRNA, with no significant effect on *Padi3* expression (**Fig. 6J**), suggesting that PAD3 protein depletion in these myoblasts is likely mediated by a post-transcriptional mechanism. Together, these findings support a previously unrecognized role for the cBAF subfamily of mSWI/SNF complexes in promoting *Padi* gene expression and protein levels, thereby enabling PAD function at myogenic loci.

We previously showed that knockdown of individual PAD enzymes selectively reduced protein levels of the targeted PAD without affecting expression of the other PADs (**Fig. 2**). Based on these data alone, one would predict that reduction of the amount of a single PAD enzyme would specifically impair chromatin binding of only that PAD. However, the reciprocal regulatory relationship between PAD enzymes and mSWI/SNF complexes described above leads to a different prediction. Specifically, knockdown of any PAD enzyme should reduce binding of all PADs at myogenic regulatory elements, due to the concomitant decrease in mSWI/SNF subunit expression and binding at target loci. Consistent with this model, knockdown of *Padi2*, *Padi3*, or *Padi4* resulted in reduced occupancy of all three PAD enzymes at the *MyhIIb* promoter and *Ckm* enhancer (**Fig. 7A-C**). A few gene-and PAD-specific exceptions were observed at the *myogenin*, *Mymk* and *Cav3* promoters (**Fig. 7A-C**), again suggesting locus-specific mechanisms that may modulate PAD recruitment. Overall, however, the data support the idea that knockdown of a specific *Padi* gene impacts the binding of multiple PADs at most myogenic genes. Collectively, these data reveal a reciprocal regulatory circuit in which PAD enzymes and mSWI/SNF complexes mutually reinforce each other’s expression and chromatin association to ensure robust activation of the myogenic gene program.

**Figure 7.**
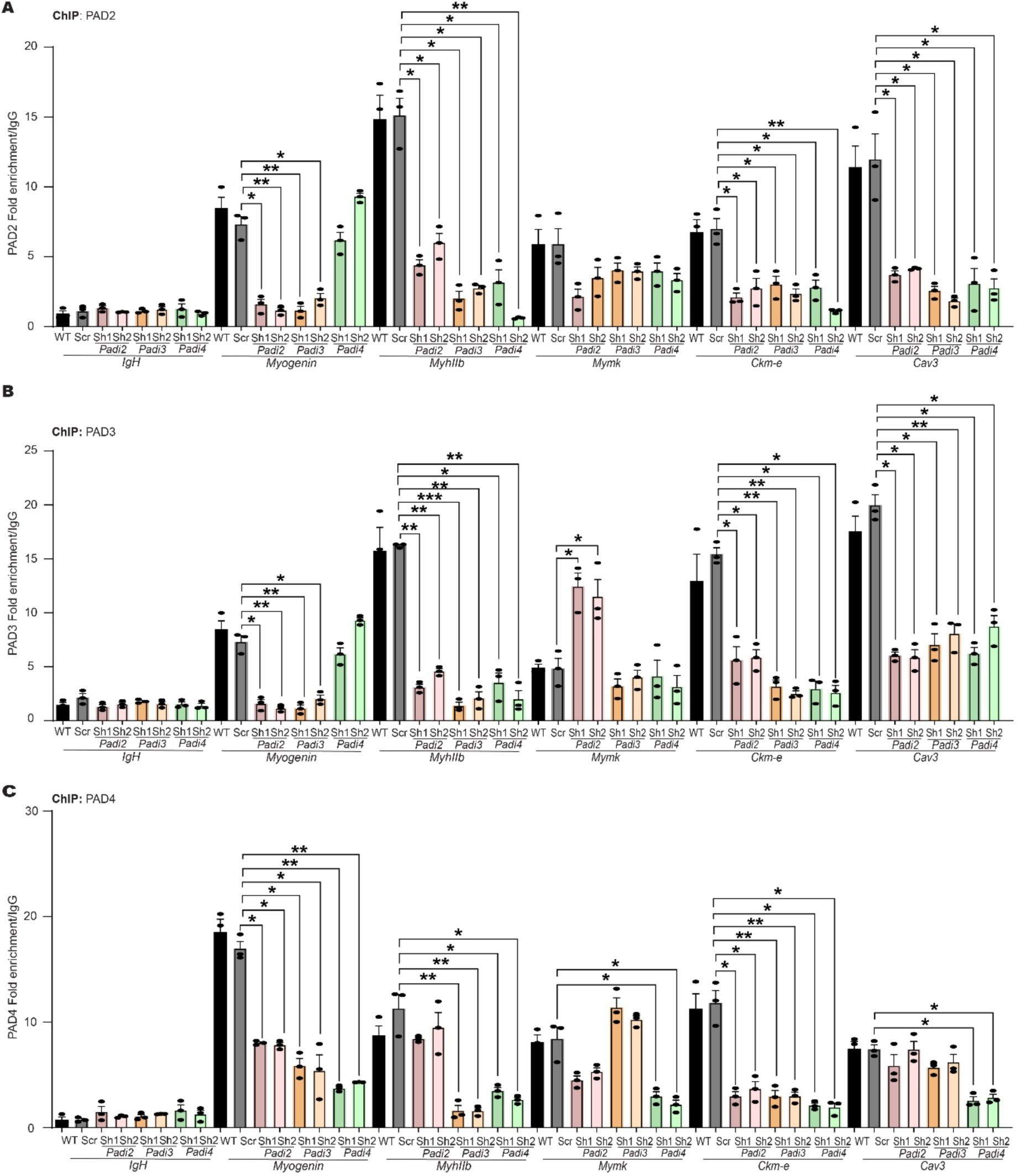
Knockdown of individual *Padi* genes decreases general PAD occupancy of myogenic gene regulatory sequences during differentiation. (A-C) ChIP-qPCR analysis showing enrichment of PAD2 (A), PAD3 (B), and PAD4 (C) at the regulatory sequences of the indicated myogenic gene in primary myoblasts differentiated for 24 h in wild-type (WT), scrambled shRNA (Scr), or *Padi2, Padi3*, or *Padi4* knockdown cells. Distinct shRNAs for each *Padi* gene are indicated by Sh1 and Sh2. Bar graphs represent mean ± SEM from three independent experiments. *p < 0.05, **p < 0.01, and ***p < 0.001.

### PAD enzymes and cBAF mSWI/SNF complexes occupy myogenic regulatory sequences in murine muscle tissue

Skeletal muscle differentiation and regeneration are governed by tightly coordinated chromatin remodeling events that couple cell state transitions to the activation of lineage-specific gene expression programs^1–5^. In adult muscle, satellite cells reside in a quiescent state beneath the basal lamina of mature myofibers and undergo rapid chromatin reorganization upon activation to support myogenic gene expression and tissue repair^34^. Building on our cell culture findings that PAD enzymes and mSWI/SNF chromatin-remodeling complexes cooperatively regulate myogenic transcription, we hypothesized that these enzymes occupy myogenic regulatory elements *in vivo* in a manner that reflects cellular differentiation state and transcriptional activity.

To test this hypothesis, tibialis anterior (TA) muscles were isolated from adult mice, and satellite cells were immediately separated from myofibers using published methods^53^. The purity and identity of each population was confirmed by qPCR analysis of established marker genes. Satellite cells were enriched for the proliferation marker *Pax7* (**Supp. Fig. 5A**), whereas myofiber preparations had elevated expression of the differentiation markers *myogenin* (**Supp. Fig. 5B**) and *MyhIIB* (**Supp. Fig. 5C**). Steady-state qPCR analysis revealed comparable expression of *Padi2*, *Padi3*, and *Padi4* in satellite cells and myofibers (**Supp. Fig. 5D-F**). Expression levels of cBAF complex subunits *Baf250a* and *Dpf2* (**Supp. Fig. 5G,H**) were also comparable in both muscle cell populations, with slightly higher levels in myofibers.

Both satellite cells and myofibers were subsequently processed for ChIP analysis^53^. PAD2, PAD3, PAD4, BAF250A, and DPF2 were detected at the *myogenin* promoter in both cell types (**Supp. Fig. 5I**). At the *MyhIIb* promoter, PAD4 was detected in both cell types. PAD2, PAD3, and the two cBAF subunits were present specifically in the myofiber samples but at or near background levels in satellite cells (compare to IgH results; **Supp. Fig. 5J,K**), suggesting that binding of these cofactors likely correlates with gene expression. Together, these data demonstrate that PAD2, PAD3, and PAD4, as well as cBAF mSWI/SNF complex subunits, occupy myogenic gene regulatory sequences *in vivo*, supporting the idea of a functional role for these enzymes in regulating myogenic transcription in skeletal muscle.

### PAD enzymatic activity is required for adult skeletal muscle regeneration

Given the in vitro requirements for PAD enzymes during myoblast differentiation and the *in vivo* occupancy of PAD enzymes at myogenic regulatory regions, we next sought to determine whether PAD activity is required for skeletal muscle regeneration following injury. We employed BB-Cl amidine, a well-characterized pan-PAD enzymatic inhibitor^54^, and AFM32a and GSK484, inhibitors of PAD2 and PAD4, respectively^25,55,56^,to assess enzyme function *in vitro* and *in vivo*.

First, we differentiated primary myoblasts in the presence of BB-Cl amidine and showed a dose-dependent inhibition of myogenic differentiation (**Fig. 8A**). This finding complements our PAD knockdown studies (**Fig. 2**) and reinforces the conclusion that PAD enzymes are required for efficient myoblast differentiation. Molecular analysis revealed a marked reduction in citrullinated H3R26, a known PAD-dependent histone modification, confirming effective inhibition of PAD enzymatic activity (**Fig. 8B**).

**Figure 8.**
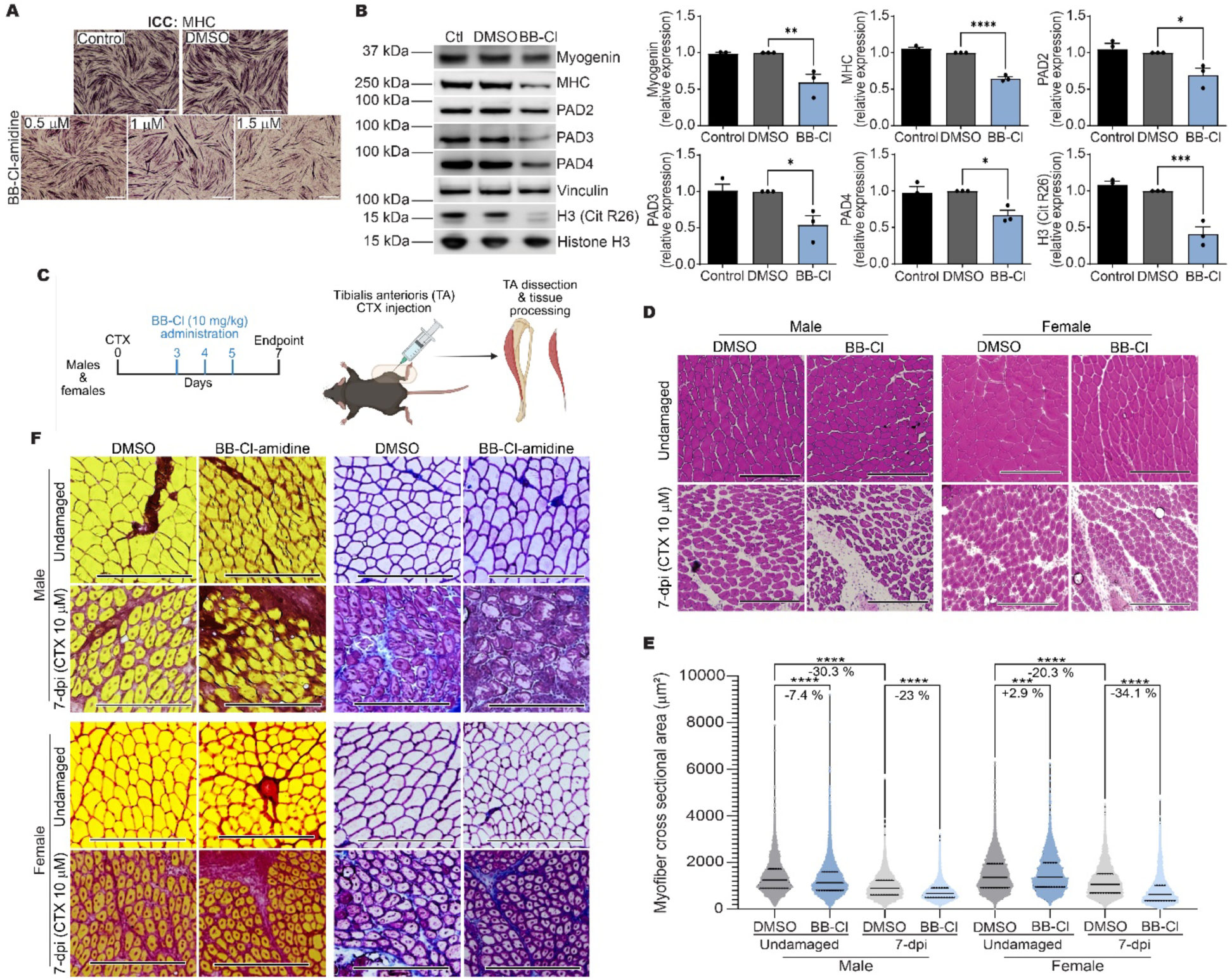
Chemical inhibition of PAD catalytic activity impairs primary myoblast differentiation and skeletal muscle regeneration. **(A)** Representative light micrographs of primary myoblasts differentiated for 24 h and immunostained for MHC under control (untreated), DMSO, or BB-Cl-amidine treatment. Scale bar, 280 μm. **(B)** Representative immunoblots (left) and quantification (right) of myogenin, MHC, PAD2, PAD3, PAD4, and the histone H3 citrullination mark, H3CitR26, in primary myoblasts differentiated for 24 h in the presence of BB-Cl-amidine (BB-Cl; 1.5 µM). **(C)** Schematic of the *in vivo* experimental design to assess the effect of BB-Cl-amidine on skeletal muscle regeneration. Cardiotoxin (CTX) was injected into the tibialis anterior (TA) muscle, followed by intraperitoneal administration of BB-Cl-amidine at 3-, 4-, and 5-days post-injury (dpi), with analysis at 7 dpi in male and female mice. Drawing created with Biorender. **(D)** Hematoxylin & Eosin (H&E) staining of representative 10 μm cross-sections of contralateral undamaged TA muscle and injured TA at 7 dpi under vehicle or BB-Cl-amidine treatment. Scale bars, 270 μm. **(E)** Quantification of myofiber regeneration by measuring fiber diameter in TA muscle at 7 dpi. **(F)** Representative PicroSirius Red and Masson’s trichrome staining of 10 μm TA muscle cross-sections from contralateral undamaged and injured muscles under vehicle or BB-Cl-amidine treatment, showing fibrosis and extracellular matrix deposition. Data in bar graphs represent mean ± SEM, N = 3; ***p < 0.001 and ****p < 0.0001.

Consistent with impaired differentiation, expression of myogenin and MHC was also reduced (**Fig. 8B**). Notably, BB-Cl amidine treatment led to a 40-50% reduction in PAD2, PAD3, and PAD4 protein levels, indicating that this PAD inhibitor impacts not only enzymatic activity but also PAD protein abundance, suggesting both direct and indirect mechanisms of action (**Fig. 8B**).

To examine PAD function during muscle regeneration *in vivo*, male and female mice were subjected to cardiotoxin (CTX) injection in the TA muscle to induce acute muscle injury^33,57,58^ (**Fig. 8C**). The contralateral TA muscle served as an uninjured control, as previously reported^33^. Three days post-injury, BB-Cl amidine or DMSO was injected into the injured muscle, with two additional injections administered at 24 h intervals. Muscles were harvested seven days post-injury, a time point corresponding to active regeneration (**Fig. 8C**). In uninjured male mice, BB-Cl amidine treatment resulted in a modest (7.4%) reduction in average myofiber cross-sectional area, as detected by H&E staining (**Fig. 8D,E**). By contrast, BB-Cl amidine treatment of CTX-injured male muscle caused a pronounced 23% reduction in myofiber cross-sectional area, representing a three-fold greater effect relative to uninjured tissue (**Fig. 8D,E**). Immunofluorescence of regenerating muscle in males showed a reduced amount of MHC in the presence of BB-Cl amidine, further supporting the idea of the pan-PAD inhibitor impairing muscle regeneration (**Supp. Fig. 6A,B**). In female mice, BB-Cl amidine had no measurable reduction in myofiber size in uninjured muscle but caused a 34% reduction in myofiber cross-sectional area in regenerating muscle compared to DMSO-treated controls (**Fig. 8D,E**). Although the basis for this sex-specific difference is not understood, these results clearly demonstrate that PAD activity is required for efficient skeletal muscle regeneration following injury.

Histological examination of regenerating muscle treated with BB-Cl amidine revealed regions devoid of myofibers (**Fig. 8D**), suggestive of increased fibrosis. Consistent with this interpretation, Sirius Red and Masson’s Trichrome staining revealed significantly increased collagen deposition in BB-Cl amidine–treated muscles from both male and female mice (**Fig. 8F**).

### PAD4 is a key determinant of efficient skeletal muscle regeneration

To dissect the contributions of individual PAD enzymes to *in vivo* muscle regeneration, we employed isoform-selective PAD inhibitors. AFM32a has been reported as a PAD2-selective inhibitor^55^, whereas GSK484 selectively inhibits PAD4^56^. No specific inhibitors of PAD3 are currently available.

Similar to our findings with *Padi* knockdown experiments, both AFM32a and GSK484 impaired the differentiation of primary myoblasts in culture (**Fig. 9A**). However, AFM32a inhibited differentiation at sub-micromolar concentrations, raising potential concerns regarding specificity. Molecular analyses confirmed reduced histone citrullination at H3R2/R8/R17, indicating effective PAD inhibition (**Fig. 9B,C**), and showed decreased expression of myogenin and MHC, consistent with impaired differentiation (**Fig. 9A**).

**Figure 9.**
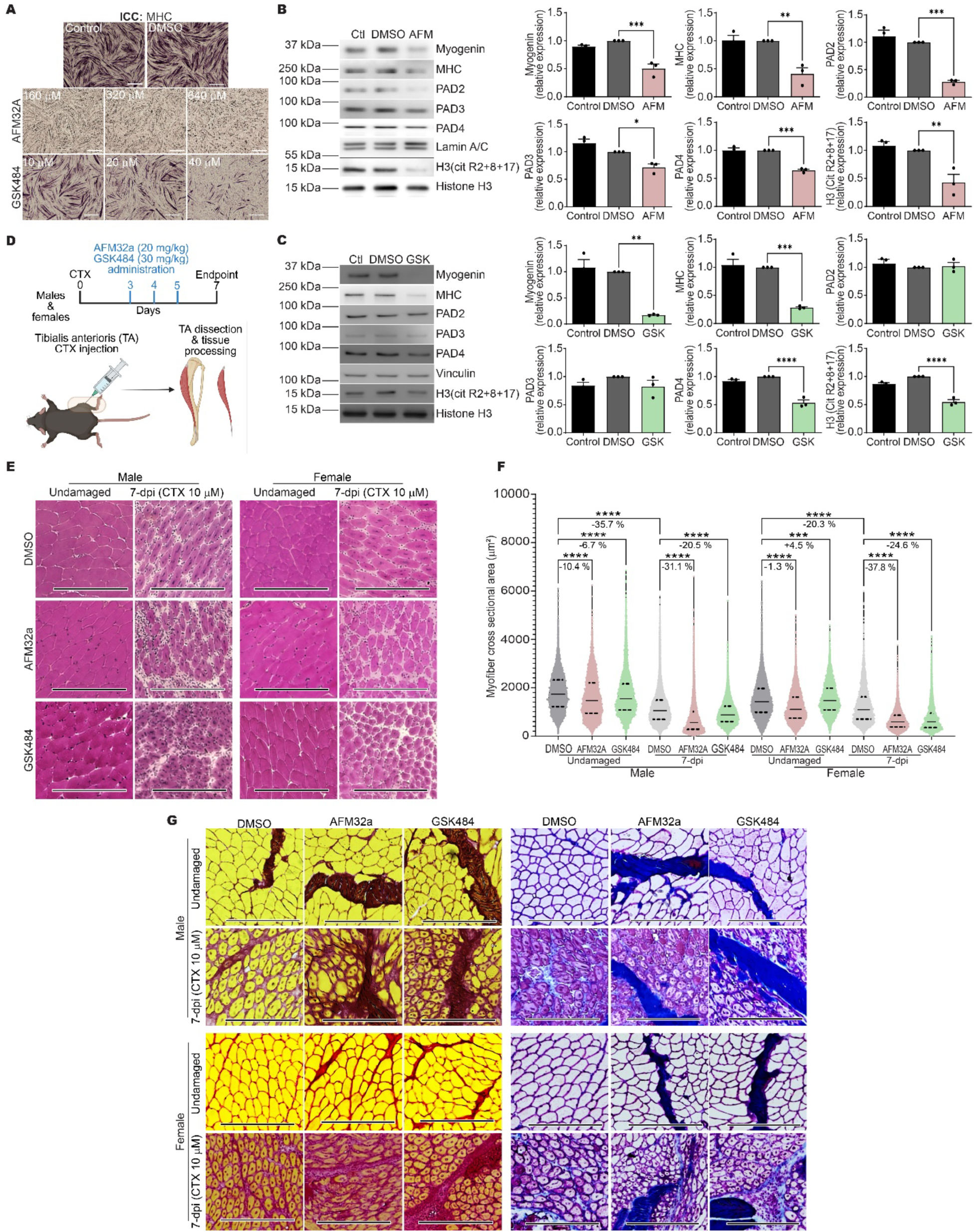
Chemical inhibition of PAD2 and PAD4 impairs myoblast differentiation and skeletal muscle regeneration. **(A)** Representative light micrographs of primary myoblasts differentiated for 24 h and immunostained for MHC under control (untreated), DMSO, AFM32A, or GSK484 treatment. Scale bar, 280 μm. **(B,C)** Representative immunoblots (left) and corresponding quantification (right) of myogenin, MHC, PAD2, PAD3, PAD4, and the histone H3 citrullination mark, H3CitR2+R8+R17, in primary myoblasts differentiated for 24 h in the presence of AFM32A (640 µM; **B**) or GSK484 (40 µM; **C**). **(D)** Schematic representation of the *in vivo* experimental design to assess the effects of PAD inhibition on skeletal muscle regeneration. Cardiotoxin (CTX) was injected into the tibialis anterior (TA) muscle, followed by intraperitoneal administration of AFM32A or GSK484 at 3-, 4-, and 5-days post-injury (dpi), with analysis at 7 dpi in male and female mice. (**E**) Hematoxylin & Eosin (H&E) staining of representative 10 μm cross-sections of contralateral undamaged TA muscle and injured TA at 7 dpi under vehicle control, AFM32A, or GSK484 treatment. **(F)** Myofiber regeneration was quantified by measuring fiber diameter. Scale bars, 270 μm. **(G)** Representative PicroSirius Red and Masson’s trichrome staining of 10 μm TA muscle cross-sections from contralateral undamaged and injured muscles under vehicle, AFM32A, or GSK484 treatment. Bar graphs represent mean ± SEM from three independent experiments (N = 3). *p < 0.05, **p < 0.01, ***p < 0.001, and ****p < 0.0001.

Notably, however, AFM32a treatment reduced the protein levels of PAD2, PAD3, and PAD4 (**Fig. 9B**), similar to what was observed with BB-Cl amidine, suggesting indirect effects on PAD expression. By contrast, GSK484 selectively reduced PAD4 protein levels without affecting the levels of PAD2 or PAD3 (**Fig. 9C**), supporting its specificity for PAD4 activity and expression.

*In vivo*, injection of AFM32a or GSK484 into regenerating muscle (**Fig. 9D**) largely phenocopied the effects observed with BB-Cl amidine (**Fig. 8**). In male mice, both inhibitors caused modest reductions in myofiber cross-sectional area in uninjured muscle, with substantially larger reductions observed in injured, regenerating tissue (**Fig. 9E,F**). Immunofluorescence studies showed that both inhibitors caused a decrease in the amount of MHC in regenerating male skeletal muscle (**Supp. Fig. 6A,C,D**). In female mice, neither inhibitor affected uninjured muscle, but both significantly impaired regeneration following injury (**Fig. 9E,F**). Increased fibrosis was also observed in inhibitor-treated muscles, as revealed by Sirius Red and Masson’s Trichrome staining (**Fig. 9G**). While interpretation of the effects of AFM32a might be confounded by indirect suppression of multiple PAD isoforms, the specificity of GSK484 strongly supports a critical role for PAD4 in skeletal muscle regeneration. Together, these data identify PAD4 as a key enzymatic regulator of myogenic differentiation and adult muscle regeneration *in vivo*.

## DISCUSSION

Here, we identified the PADs as essential and previously underappreciated regulators of skeletal muscle differentiation and regeneration. We showed that PAD2, PAD3, and PAD4 are induced during myoblast differentiation, localize to the nucleus, bind myogenic gene regulatory regions, and are required for efficient activation of myogenic gene expression and myoblast differentiation *in vitro* and for adult muscle regeneration *in vivo*. Notably, we report a reciprocal functional dependency between PAD enzymes and cBAF-type mSWI/SNF chromatin remodeling complexes, revealing a chromatin regulatory circuit that promotes myogenic gene activation.

All three PAD enzymes contributed to myoblast differentiation, but with partially distinct transcriptional consequences. *Padi2* and *Padi4* knockdown broadly impaired myogenic and metabolic gene expression programs, whereas *Padi3* KD reduced differentiation despite comparatively modest effects on canonical myogenic transcripts. Instead, PAD3 depletion altered core cellular metabolic processes, suggesting that PAD3 supports myogenesis indirectly by maintaining cellular states permissive for differentiation. These findings point to functional specialization among PAD family members, even as they converge on a shared tissue differentiation outcome.

Our results may or may not contrast with data generated via mouse gene knockout studies. *Padi2* and *Padi4* single and double knockouts exist with assorted non-lethal, non-skeletal muscle phenotypes^59^, as do *Padi3* knockout mice^60^. The obvious conclusion is that there is redundancy among the PADs during embryonic development. However, our *in vitro* studies utilized primary myoblasts derived from satellite cells, which are adult stem cells, and our *in vivo* investigation of muscle regeneration after injury evaluates adult myogenesis. To our knowledge, none of those genetically altered mouse models have been tested for muscle regeneration capacity. Therefore, our results demonstrating specific, non-redundant requirements for PAD2, PAD3, and PAD4 in adult myogenesis are not incompatible with prior findings.

Consistent with a direct role in transcriptional regulation, PAD2, PAD3, and PAD4 dynamically associated with regulatory regions of active myogenic genes, while dissociating from the *Pax7* promoter as differentiation progressed. This pattern closely mirrored transcriptional activity, indicating that PAD-chromatin interactions are regulated during lineage commitment. PAD binding to target genes also generally correlated with histone H3 citrullination at R26 and when R2/R8/R17 were examined together.

Interestingly R17 citrullination by itself did not correlate with transcriptional activity, suggesting the correlating citrullination function occurs at R2 and/or R8. However, knockdown of individual PAD enzymes did not yield residue-specific or gene-independent effects on histone citrullination, indicating redundancy, context dependence, or contributions from non-histone substrates. Thus, histone citrullination likely represents only one component of a broader PAD-dependent regulatory program.

Our *in vivo* studies indicate a role for PAD enzymes in normal adult skeletal muscle regeneration. Pharmacological inhibition of PAD activity impaired regeneration following injury and increased fibrosis, demonstrating that PAD enzymes are required not only for myoblast differentiation but also for proper tissue remodeling during repair. Isoform-selective inhibition identified PAD4 as a critical contributor to this process, as PAD4 inhibition alone was sufficient to compromise regeneration and promote fibrosis without broadly suppressing other PAD family members. We believe it is likely that the PAD2 isoform also plays a critical role in regeneration, but the likely indirect effect of the PAD2 inhibitor on the protein levels of PAD3 and PAD4 prevents us from making such a conclusion.

A central and unexpected finding of this study is the discovery of reciprocal regulation between PAD enzymes and subunits of the cBAF subfamily of mSWI/SNF chromatin remodelers. PAD knockdown reduced chromatin binding of cBAF-specific subunits BAF250A and DPF2 and decreased their expression, while knockdown of these cBAF subunits diminished PAD binding to chromatin and reduced PAD expression. This bidirectional dependency provides a mechanistic explanation for the observation that KD of any single PAD enzyme impaired chromatin association of all PAD family members. Our data supports a model in which PAD enzymes and cBAF chromatin remodelers form a mutually reinforcing regulatory circuit that stabilizes chromatin for myogenic gene activation. This observation has implications for current efforts seeking to target PAD and mSWI/SNF enzymes in human disease. Mis-regulation of PAD enzymes are implicated in autoimmune disease and cancer and mis-regulation of mSWI/SNF enzymes is implicated in many cancers as well as in neurodevelopmental diseases^26–29^.

Significant effort and resources have been invested in the development of small molecule inhibitors, proteolysis targeting chimeras (PROTACs), and biologic approaches to modifying disease-related PAD and mSWI/SNF protein functions^22,61–63^. Our work introduces the idea that therapeutic approaches to modifying PAD expression and function could be applied to cancers and neurodevelopment diseases associated with mSWI/SNF subunit mis-regulation and that therapeutic approaches to modifying mSWI/SNF subunit expression and function might impact autoimmune diseases and cancers associated with PADs.

In conclusion, our work provides a foundation to further investigate basic mechanisms by which PAD enzymes integrate into chromatin regulatory networks to influence developmental and regenerative outcomes in skeletal muscle as well as explore translational approaches to multiple human diseases.

## MATERIALS AND METHODS

### Antibodies

Hybridoma supernatants against myosin heavy chain (MHC) and myogenin (MF20 and F5D, deposited by D. A. Fischman and W.E. Wright, respectively) were obtained from the Developmental Studies Hybridoma Bank (University of Iowa).

Mouse anti-BRG1(G-7; sc-17796), anti-PAD3 (F6; sc-393622), anti-PAD4 (A11; sc-365369), anti-lamin A/C (4A58), and anti-α-tubulin (B-7; sc-5286) were obtained from Santa Cruz Biotechnologies. Rabbit anti-BAF250A (A18650) was from Abclonal Technologies Rabbit anti-PAD2 (E3P8Z; #97647), anti-vinculin (E1E9V; #13901), anti-Histone H3 (#9715), Histone H3Cit R17 (E4O3F; #97272), and anti-DPF2 (E7N8J; #71642) were from Cell Signaling Technologies. Anti-Histone H3Cit R2+R8+R17 (ab5103) and anti-Histone H3Cit R26 (EPR20606; ab212082) were from Abcam. The secondary antibodies used for western blots were the HRP-conjugated anti-mouse and anti-rabbit (31430 and 31460, respectively) and for immunofluorescence were the goat anti-rabbit IgG Alexa Fluor Plus 488 and 633 (A11008, A21070) and the goat anti-mouse IgG Alexa Fluor Plus 488 (A11001) from Thermo Fisher Scientific.

### Cell culture

Murine primary myoblasts were obtained from iXCells Biotechnologies (10MU-033). Proliferating myoblasts were maintained in growth medium consisting of 1:1 (v/v) DMEM:F-12 (11039-021; Life Technologies), 20% fetal bovine serum (PS-100; Phoenix Scientific), 25 ng/ml basic fibroblast growth factor (FGF; 4114-TC, R&D Systems, Inc.) and 1% antibiotics (penicillin/streptomycin; 151-40-122, Gibco).

Proliferating C2C12 myoblasts were obtained from the ATCC (CRL-1772) and were maintained at subconfluent densities in proliferation medium containing Dulbecco’s modified Eagle’s medium (DMEM) supplemented with 10% fetal bovine serum (FBS) and 1% penicillin/streptomycin.

To induce myoblast differentiation, primary myoblasts were seeded on plates coated with 1% Matrigel (354234; Corning, Inc.). Then, primary myoblasts and C2C12 cells were allowed to reach 90% confluency, and differentiation was initiated by replacing the proliferation medium with differentiation medium composed of DMEM (11965-092; Life Technologies), 2% horse serum (16050-122; Life Technologies), 1% antibiotics, and insulin/transferrin/selenium (51300-044; Gibco).

The HEK293T cell line (CRL-3216), obtained from ATCC, is a derivative of human embryonic kidney cells that expresses a mutant SV40 large T antigen and was used for lentiviral production. HEK293T cells were maintained at sub-confluent densities in proliferation medium consisting of DMEM supplemented with 10% FBS and 1% penicillin/streptomycin. All cultures were maintained in a humidified incubator at 37°C with 5% CO_2_.

### Lentivirus production for stable shRNA transduction in primary and C2C12 myoblasts

Mission plasmids (Sigma-Aldrich) encoding a scrambled control shRNA (Scr) or two distinct shRNAs targeting *Padi2*, *Padi3*, or *Padi4* **(Supp. Table 2)** were purified using the PureYield Plasmid Midiprep System (Zymo Research) according to the manufacturer’s instructions. For lentivirus production, each shRNA plasmid (15 µg) together with the packaging vectors pLP1 (15 µg), pLP2 (6 µg), and pSVGV (3 µg), were transfected into HEK293T cells using Lipofectamine 2000 (Thermo Fisher Scientific). Viral supernatants were collected at 24 and 48 h post-transfection, then filtered through a 0.22 µm syringe filter (Millipore). Proliferating primary and C2C12 myoblasts were transduced in the presence of 8 µg/mL polybrene and selected with 2 µg/mL puromycin (A11138-03; Gibco) as previously reported ^51,64–69^. Cells were maintained with 1 μg/mL puromycin.

### Myoblast treatment with PAD enzyme inhibitors

Primary myoblasts were cultured under differentiation conditions with the pan-PAD enzyme inhibitor BB-Cl amidine (HY-111347; MedChemExpress), the PAD2 inhibitor, AFM32A synthesized as previously described^55^, and the PAD4 inhibitor GSK484^25,56^ (SML1658; MilliporeSigma). To determine the optimal working concentrations, a range of inhibitor concentrations was titrated. BB-Cl amidine was tested at 0.5-1.5 µM, AFM32A at 180-640 µM, and GSK484 at 10-40 µM. The inhibitor-containing medium was refreshed every 24 h. Final working concentrations are indicated in the corresponding figure legends.

### Subcellular fractionation

Cell fractionation was carried out using the rapid, efficient, and practical (REAP) method ^70^. Briefly, differentiated primary and C2C12 myoblasts were washed three times with ice-cold 1× PBS and resuspended in 1 mL of PBS supplemented with a protease inhibitor cocktail (PIC; PIA32955; Thermo Fisher Scientific). After centrifugation, the supernatant was discarded, and the cell pellet resuspended in 900 μL of ice-cold PBS containing 0.1% NP-40. Cells were pipetted 30-40 times with a P1000 micropipette tip to disrupt membranes and release nuclei. The integrity of isolated nuclei was verified by light microscopy. An aliquot of the suspension (300 μL) was saved as the whole-cell lysate, and the remaining fraction was centrifuged at 10,000 rpm. The supernatant (cytosolic fraction) was transferred to a new tube, while the nuclear pellet was washed once with 1 mL of PBS containing 0.1% NP-40. The nuclear fraction was centrifuged again at 10,000 rpm, the supernatant discarded, and the pellet resuspended in 150 μL of PBS. Fractions were analyzed by WB using antibodies against PAD2, PAD3, and PAD4, with lamin A/C and α-tubulin used as controls. Signals were visualized with the Tanon chemiluminescence detection system (180-5001; Abclonal Technologies).

### Western blot analyses

Primary and C2C12 myoblasts, as well as control and knockdown cells, were washed three times with ice-cold PBS and solubilized with RIPA buffer [10 mM piperazine-N,N-bis(2-ethanesulfonic acid), pH 7.4; 150 mM NaCl; 2 mM ethylenediaminetetraacetic acid (EDTA); 1% Triton X-100; 0.5% sodium deoxycholate; and 10% glycerol] containing PIC. Protein concentration was determined by Bradford ^71^. Twenty micrograms of each sample were prepared for SDS-PAGE by boiling in Laemmli buffer. The resolved proteins were electro-transferred to PVDF membranes (IPVH00010; Millipore). Proteins of interest were detected with specific antibodies as indicated in the figure legends, followed by species-specific peroxidase-conjugated secondary antibodies and visualization using the Tanon chemiluminescence detection system (180-5001; Abclonal Technologies).

### Chromatin immunoprecipitation assays

Chromatin immunoprecipitation (ChIP) assays were conducted on primary myoblasts in triplicate from three independent biological replicates as previously described ^51,65,68,72,73^. Briefly, crosslinked lysates from myoblasts were incubated overnight with antibodies targeting PAD2, PAD3, PAD4, BAF250A, DPF2, or with normal rabbit IgG as a control. Following reversal of crosslinking, DNA was purified using ChIP DNA Clean and Concentrator columns (D5201; Zymo Research) according to the manufacturer’s protocol. Quantitative PCR was performed using Fast SYBR Green Master Mix (4385612; Thermo Fisher Scientific) on an ABI StepOne Plus Sequence Detection System (Applied Biosystems). Primer sequences are provided in **Supp. Table 3**. Data was analyzed using the comparative Ct method (ΔCt; ^74^) to determine the percentage of total input DNA immunoprecipitated by each antibody.

### Steady state gene expression analysis

Total mRNA was obtained from three independent biological replicates of controls, knockdown and inhibitor-treated primary and C2C12 myoblasts with TRIzol (Invitrogen) following the manufacturer’s instructions. cDNA synthesis was performed with 900 ng of RNA as template; for the reverse transcription reaction, we used HiFiScript gDNA Removal RT MasterMix Kit (CW2020M; Cwbio) following the manufacturer’s protocol. Quantitative RT-PCR was performed with Fast SYBR green master mix on the ABI StepOne Plus Sequence Detection System using the primers listed in **Supp. Table 3**, and the delta threshold cycle value (ΔCt; ^74^) was calculated for each gene and represented the difference between the Ct value of the gene of interest and that of the control gene, *Eef1a1*.

### Immunocytochemistry

Control, KD and inhibitor-treated primary and C2C12 myoblasts were fixed overnight in 10% formalin in PBS at 4°C. Cells were washed three times with PBS and permeabilized for 30 min in PBS containing 0.2% Triton X-100.

Immunocytochemistry was performed on differentiating cells using myogenin and MHC hybridoma supernatants (FD5 and MF20), respectively. Samples were developed with the Universal ABC kit and Vector VIP substrate Kit, peroxidase (PK-6200, SK-4600; Vector Labs) according to the manufacturer’s instructions. Images were acquired using an Echo Rebel microscope with a 20X objective (Echo).

### Immunofluorescence in cultured myoblasts and imaging

Differentiating primary myoblasts (24 h) were grown on coverslips and fixed overnight in methanol at 4°C. The monolayers were washed with PBS, permeabilized with 0.2% Triton X-100 in PBS for 15 min and incubated for 1 h at room temperature in blocking solution containing PBS, 0.2% Triton X-100, and 3% FBS. Samples were then incubated overnight at 4°C with anti-PAD2, anti-PAD3, or anti-PAD4 antibodies (1:100 dilution) in blocking buffer. Then, the cells washed in PBS-Tween 20 (0.5%) and incubated for 3 h at room temperature with a goat anti-rabbit Alexa 633 or goat anti-mouse Alexa 488 secondary antibodies in blocking solution. Nuclei were stained by a 30 min incubation with DAPI (F6057, Sigma Aldrich). Microscopy and image processing were performed using a Leica SP8 confocal microscope, and image analyses was performed with Leica Lite software.

### Animal Care and Use Statement

All animal procedures and experimental protocols were conducted in compliance with relevant ethical regulations and were approved by the Institutional Animal Care and Use Committees (IACUCs) at the University of Wisconsin-Madison and at Wesleyan University. Experiments were performed in C57BL/6 male and female adult mice (7-8 weeks; Jackson Laboratory).

### Isolation and processing of satellite cells and myofibers from mice

Satellite cells and myofibers were as previously described^53^. Briefly, after dissection of skeletal muscle from both hindlimbs, satellite cells and myofibers were separated. Minced skeletal muscle tissue was partially digested by resuspension in 100 U/mL collagenase type II in PBS supplemented with 1 mM CaCl_2_. Samples were incubated with agitation at 37°C for 1 h in a temperature-controlled rotator. Following collagenase treatment, satellite cells were separated from mature myofibers by gravity filtration through a 70-μm nylon cell strainer into a 50 mL conical tube at room temperature. The flow-through fraction was enriched for satellite cells. To isolate the fraction retained on the membrane (myofibers), the filtration unit was inverted and gently tapped over a 50 mL conical tube to facilitate transfer of the myofibers into the tube. Samples were placed on ice and centrifuged at 300 × *g* at 4°C for 5 min and the supernatant was removed by aspiration or pipetting. A portion of each sample was used for RNA analysis; small aliquots of tissue were transferred into 1.5-mL microcentrifuge tubes, and mRNA extraction was performed as described above.

For ChIP assays, the pellets were resuspended in seven volumes of lysis buffer (10 mM HEPES-KOH, pH 7.3; 10 mM KCl; 5 mM MgCl_2_; 0.5 mM DTT) containing freshly added protease inhibitors (0.2 mM PMSF and 10 μg/mL leupeptin). For the myofiber fraction, the lysis buffer was supplemented with 3 μg/mL cytochalasin B. Samples were incubated on ice for 30 min. Large fragments of tissue were disrupted using a tissue homogenizer, resulting in a homogenous cell suspension. Samples were centrifuged at 3,000 x *g* for 5 min at 4°C and pellets were resuspended in 2.5 volumes of 10-STM buffer (10% sucrose, 10 mM triethanolamine pH 7.5, 5 mM MgCl_2_, 10 μg/mL leupeptin).

Twice the original pellet volume of 2.0 M sucrose in 10 mM Tris-HCl (pH 7.4) and 5 mM MgCl_2_ was added, and samples were gently mixed by pipetting. The suspension was transferred to a prechilled 10-mL Dounce homogenizer and further homogenized with seven slow strokes, moving the pestle carefully to minimize foaming and preserve nuclear integrity. Nuclei release and integrity were confirmed by light microscopy following staining of a small aliquot (1–2 μL).

For ultracentrifugation, 750 μL of 2.0 M sucrose buffer (10 mM Tris–HCl pH 7.4, 5 mM MgCl_2_) was added to the bottom of a 5-mL ultracentrifuge tube, and the Dounce-homogenized nuclear suspension was gently overlaid. Samples were ultracentrifuged at 116,100 × g for 1 h at 4°C. The supernatant was carefully removed, and the resulting nuclear pellet was resuspended in 500 μL of lysis buffer supplemented with 0.1% NP-40. Cross-linking was performed by adding formaldehyde to a final concentration of 1% and incubating for 5 min at room temperature, followed by quenching with glycine to a final concentration of 0.125 M for 5 min at room temperature. Cross-linked nuclei were pelleted by centrifugation at 13,500 × g for 1 min at room temperature, and the supernatant was discarded. Pellets were either flash-frozen in liquid nitrogen and stored at-70 °C or thawed on ice for immediate use. For ChIP, cross-linked nuclei were resuspended in 400 μL of lysis buffer, and the assay was performed as described above.

### Cardiotoxin muscle injury experiments

Injury experiments were performed as previously described^75^ using C57BL/6 male and female adult mice (7-8 weeks; Jackson Laboratory). Prior to initiating the procedure, the isoflurane anesthetic station was inspected to confirm proper operation. For anesthesia induction, the vaporizer was set to 4-5% (v/v) isoflurane and reduced to 2% (v/v) for maintenance during surgery. Mice were weighed, assessed for overall wellness, and placed into the anesthetic chamber with oxygen flow maintained at 2 L/min. To prevent anesthetic leakage, the connection to the nose cone was closed during induction. Once anesthetized, animals were transferred to the nose cone with isoflurane delivery sustained at 2% (v/v). Meloxicam was used an analgesic at a dose of 5 mg/kg and was administered sub-cutaneous. For surgical preparation, the mouse was placed supine, and hair on the hind limbs was removed with an electric trimmer to expose the tibialis anterior (TA) tendon. Depth of anesthesia was verified by the absence of a pedal withdrawal reflex^33,76^.

To induce injury, 50 µL of cardiotoxin (CTX) dissolved in saline was injected into the TA muscle using an insulin syringe^33,76^. The needle was inserted parallel to the tibia at the TA tendon-muscle junction, with CTX delivered as the needle advanced toward the knee. Following injection, mice were placed in a 37°C recovery chamber until ambulatory and then transferred to sterile recovery cages. Then, the mice received intraperitoneal injections of the inhibitors BB-Cl amidine (10 mg/kg^25^), AFM32a (20 mg/kg^55^), or GSK484 (30 mg/kg^56^) on days 3, 4, and 5 post-injury (**Figs. 8C, 9D**).

Animals were maintained for 7 days post-injury before sacrifice, at which time TA muscles were dissected and collected for downstream analyses.

### Hematoxylin and Eosin (H&E) staining

H&E staining of TA cross-sections was performed to determine differences in general morphology of TA in controls and PAD-inhibitor treated animals. Slides were rehydrated in deionized water (ddH_2_O) for 2 min, followed by incubation with Mayer’s hematoxylin for 10 min with gentle agitation. After staining, slides were rinsed in water for 10 s and immersed in saturated Li_2_CO_3_ (pH 8.5) for 1 min to enhance nuclear bluing. Sections were washed twice in water (10 s each), immersed in 70% ethanol for 1 min, and counterstained with Eosin Y (pH 5.3) for 5 min. Slides were then sequentially dehydrated with 70% ethanol (1 min), 100% ethanol (two 1-min rinses), and xylene (two 1-min rinses). Stained sections were imaged by light microscopy, and myofiber diameters were quantified using MouseLand/Cellpose software^76,77^.

### PicroSirius Red Staining

PicroSirius Red staining was used to detect the presence of collagen and fibrosis^78,79^ in TA muscles upon injury in control and animals treated with PAD inhibitors. First, the slides were rehydrated using a graded ethanol series (100%, 80%, 40%, 3 min each), followed by one wash with water (5 min), then fixed in 4% formaldehyde (30 min), and rinsed twice with water (5 min) before. The slides were stained in Picro-Sirius red solution (1% m/v) for 1 h, then were washed in two changes of acidified water (2 min). Slides were dehydrated in three changes of 100% ethanol (1 min), cleared in xylene (1 min), and mounted in Poly Mount (Xylene Based; Polysciences). Images were captured using an Echo microscope.

### Mason’s trichrome staining

Mason’s trichrome staining was used to compare fibrosis and connective tissue deposition^80,81^ in control and PAD-inhibitor treated animals. Slides were rehydrated through a graded ethanol series (100%, 80%, and 40%; 3 min each), rinsed in water (5 min), fixed in 4% formaldehyde for 30 min, and washed twice in water (5 min each). Mason’s Trichrome staining (ab150686; Abcam) was then performed according to the manufacturer’s instructions. Briefly, slides were incubated in preheated Bouin’s solution for 1 h, cooled for 10 min, and rinsed in ddH_2_O (1 min). Nuclei were stained with Weigert’s Iron Hematoxylin (5 min), followed by a 2 min water rinse. Slides were then incubated in Biebrich Scarlet/Acid Fuchsin solution for 15 min, rinsed in ddH_2_O (1 min), sections were differentiated in phosphomolybdic/ phosphotungstic acid solution (10-15 min), or until collagen was no longer red. Collagen was stained with Aniline Blue solution (10 min), rinsed in ddH_2_O (1 min), and treated with 1% Acetic Acid (5 min). Sections were dehydrated rapidly through two changes of 95% ethanol (1 min each) and two changes of absolute ethanol (1 min each), cleared in xylene (1 min), and mounted with Poly Mount (Xylene Based). Stained sections were imaged using an Echo microscope.

### Immunofluorescence of mouse skeletal muscle tissue

Immunofluorescence staining of male TA muscle cross-sections was performed as previously described^81^. The muscle cross-section slides were equilibrated to room temperature, washed with PBS, and then fixed with 4% (w/v) PFA for 30 min, followed by two washes with PBS for 5 min each. The slides were incubated in antigen retrieval solution (0.1 M citric acid, 0.1 M sodium citrate stock (pH 6.0), and 0.2% Tween 20) at 98°C for 10 min. The slides were then washed twice with TBS for 5 min each. Permeabilization was carried out with 0.1 M glycine and 0.1% Triton X-100 in TBS on a shaker for 10 min, followed by two additional TBS washes (5 min). Slides were blocked in 5% (v/v) goat serum normal IgG (1:2000) and 2% (w/v) BSA in TBST for 1 h at room temperature. The blocking solution was aspirated and replaced with primary antibody-blocking solution, and the slides were incubated overnight at 4°C. After three washes with TBST, the slides were blocked (1 h) and were incubated with secondary antibody (1 h). Finally, the slides were washed three times with TBST, twice with TBS, and once with PBS (5 min each). Immunofluorescence images were acquired using a Leica SP8 confocal microscope and analyzed with Leica lite software.

### Statistical Analysis

Statistical analyses were performed using GraphPad Prism 10.0.0, Multiple data point comparisons and statistical significance were determined using one-way analysis of variance (ANOVA) and the comparisons were performed using Dunnett‘s multiple comparison tests. Results where p< 0.05 were considered to be statistically significant.

## AUTHOR CONTRIBUTIONS

Reagent generation and/or data acquisition and/or data analysis: MO-F, TS, AP, LB, SH, TP-B. Study design: MO-F, FJD, TP-B, ANI. Manuscript drafting: TP-B, ANI, Manuscript review: all authors

## ACKNOWLEDGEMENTS

We thank members of the Dilworth and Padilla-Benavides laboratories for technical support.

