## Supplemental Figures and Tables for "Reciprocal regulation between the protein arginine deiminases and mSWI/SNF chromatin remodelers controls skeletal muscle differentiation and regeneration"

### Supplemental figure 1

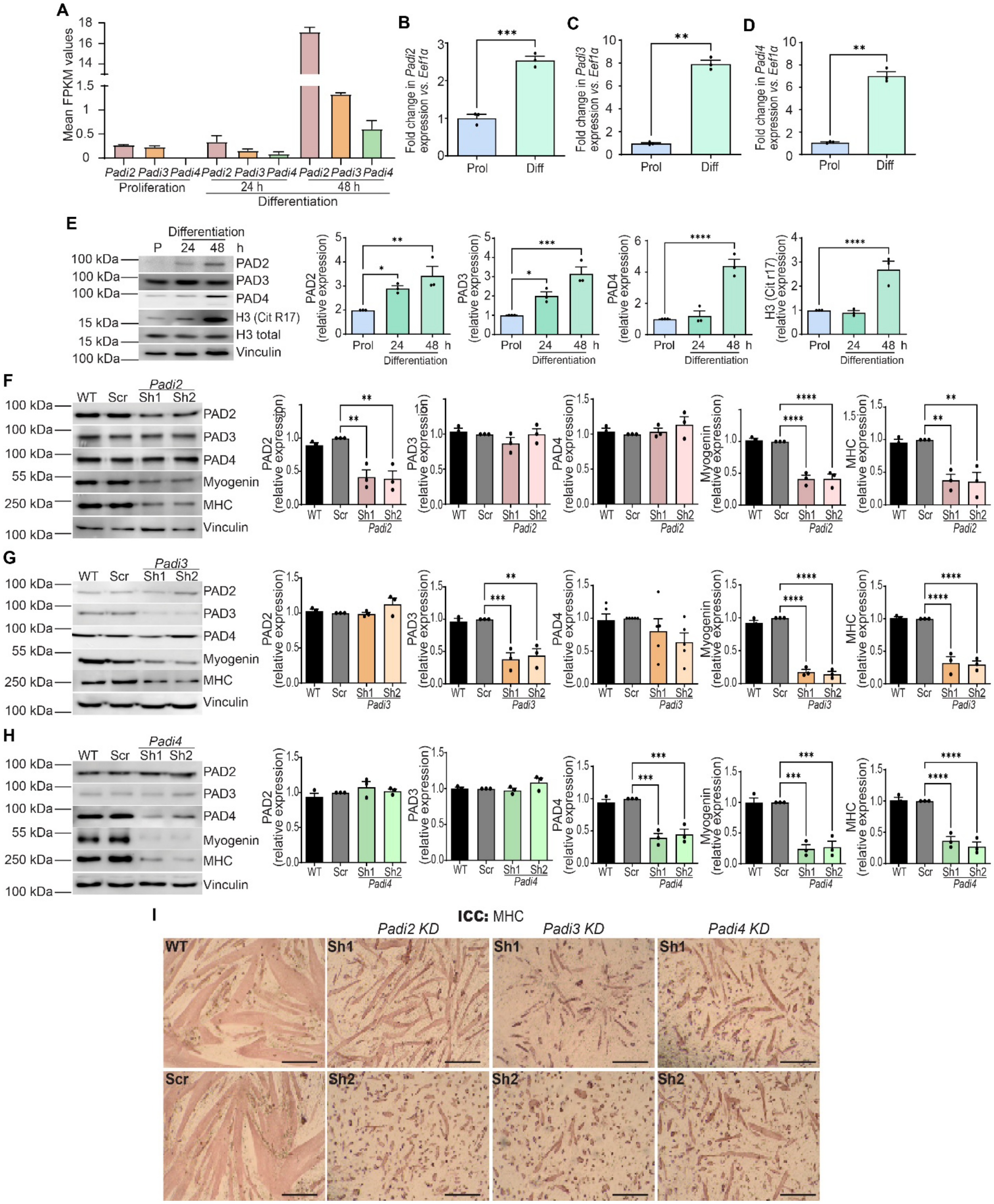

**Supplemental Figure 1: PAD2, PAD3, and PAD4 are induced during and are required for C2C12 myoblast differentiation. (A)** FPKM values for *Padi2*, *Padi3*, and *Padi4* gene expression in the RNA-seq data set (REF) from proliferating C2C12 myoblasts and from C2C12 myoblasts differentiated for 24 or 48 hours (h). **(B-D)** qPCR validation of *Padi2*, *Padi3*, and *Padi4* mRNA levels in proliferating (Prol) and 48 h differentiated (Diff) C2C12 myoblasts. **(E)** Left; Representative immunoblots showing PAD2, PAD3, PAD4, and citrullinated histone H3 (H3 CitR17) protein levels in proliferating (P) and 24 and 48 h differentiating (Diff) C2C12 myoblasts. Right: Corresponding quantification of PAD protein and citrullinated H3 levels. Total H3 was used as a normalization control for the citrullinated H3; Vinculin was used as a normalization control for the PADs. **(F-H)** Left; Representative immunoblots showing PAD2, PAD3, or PAD4, respectively, as well as myogenin, MHC and vinculin (as a control) in 48 h differentiated Wild type (WT) C2C12 myoblasts or in C2C12 myoblasts expressing scrambled sequence (Scr), *Padi2*, *Padi3*, or *Padi4* shRNA. Two distinct shRNAs (indicated as Sh1 and Sh2) were used for *Padi* gene knockdowns. Right: Corresponding quantification of PAD and myogenic marker proteins Vinculin was used as a normalization control. Data are presented as mean ± SEM from three independent experiments with $*p < 0.05$, $**p < 0.01$, $***p < 0.001$, ****p , 0.0001. **(I)** Immunocytochemistry (ICC) of the indicated differentiated C2C12 cells stained for MHC. KD, knockdown.

**Supplementary figure 2**

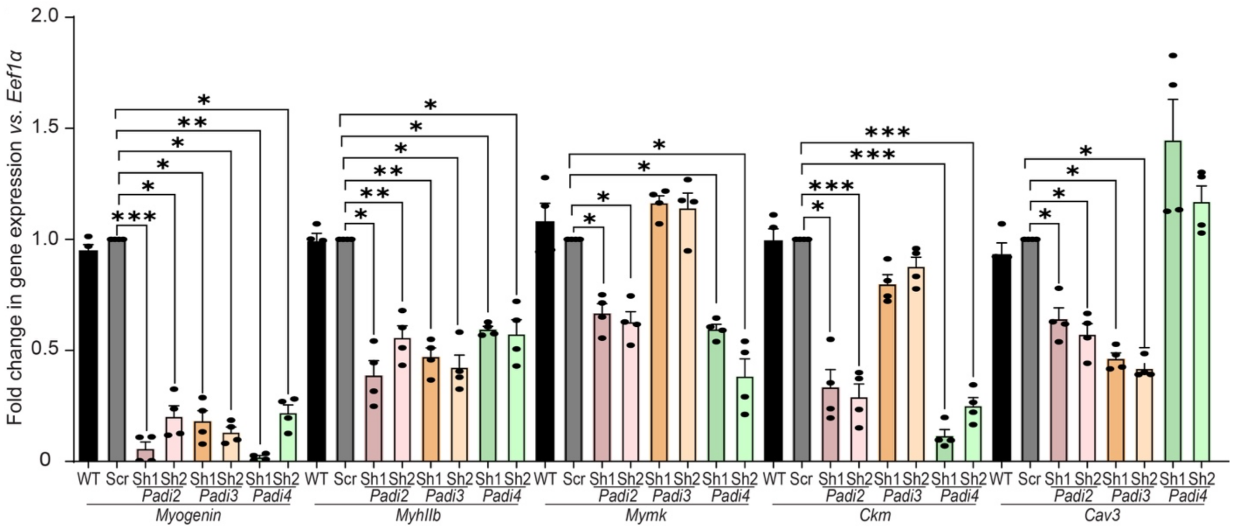


**Supp Figure 2. *Padi2, Padi3,* or *Padi4* knockdown disrupts expression of myogenic genes during differentiation.** Relative mRNA levels of specific myogenic differentiation genes (*Myogenin, MyhcIIb, Mymkr, Ckm,* and *Cav3*) were quantified by qPCR in differentiating WT, Scr, *Padi2*, *Padi3*, or *Padi4* knockdown primary myoblasts.

### Supplemental figure 3

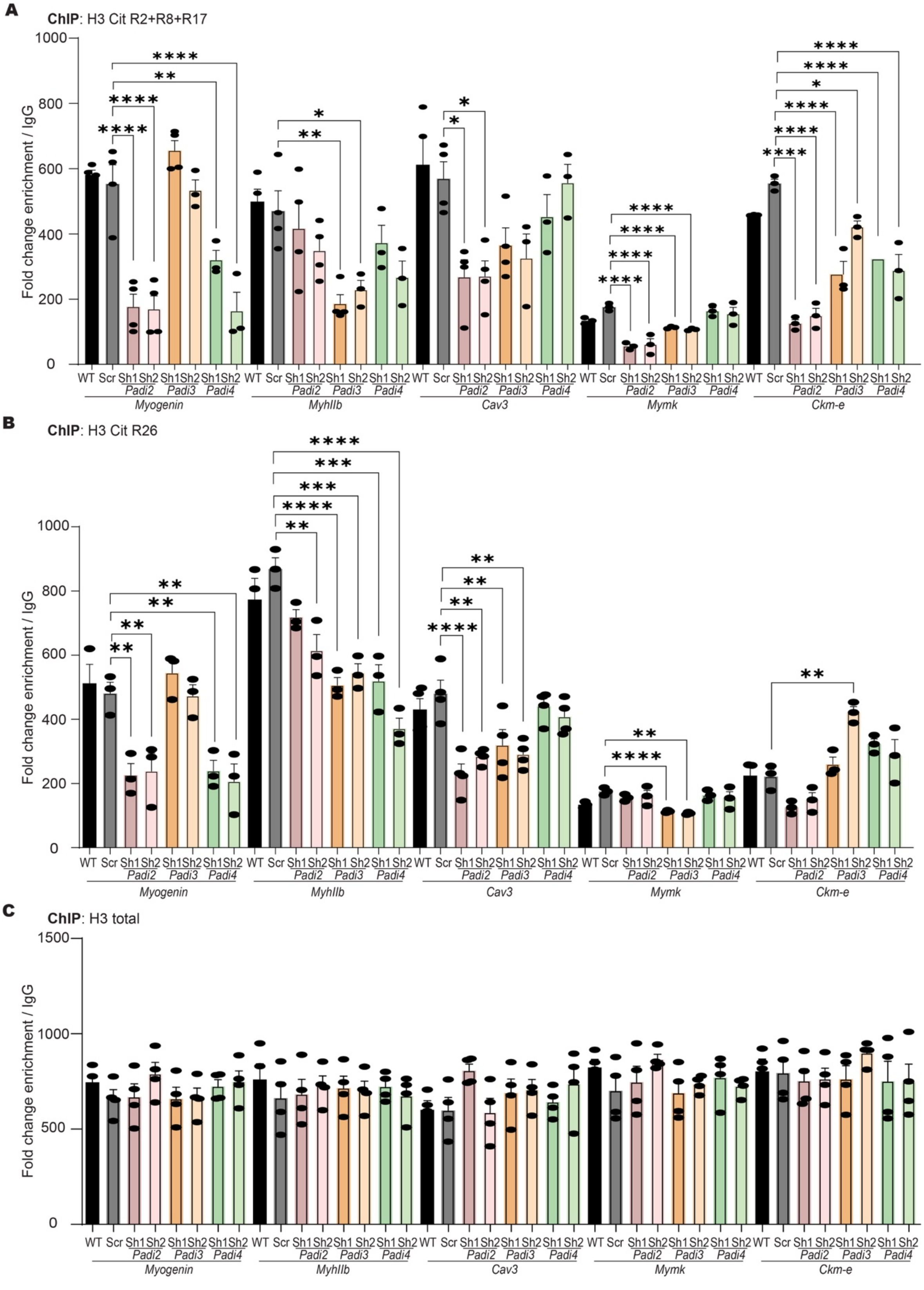

**Supplemental Figure 3: Incorporation of H3 (cit R2/R8/R17) and H3 (cit R26) at myogenic gene regulatory sequences is variably affected by knockdown of *Padi2*, *Padi3*, or *Padi4*.**

**(A-C)** ChIP experiments for H3 (cit R2/R8/R17), H3 (cit R26), and total H3, respectively, at the indicated myogenic gene regulatory sequences in primary myoblasts differentiated for 24 h following shRNA-mediated knockdown of *Padi2, Padi3*, or *Padi4*. Distinct shRNAs targeting each *Padi* gene are indicated by Sh1 and Sh2. The IgH enhancer was included as a negative sequence control. Bar graphs represent the mean ± SEM from three independent experiments. *$p < 0.05$, **$p < 0.01$, ***$p < 0.001$, and ****$p < 0.0001$. WT, Wild-type; Scr, scrambled sequence shRNA.

### Supplemental figure 4

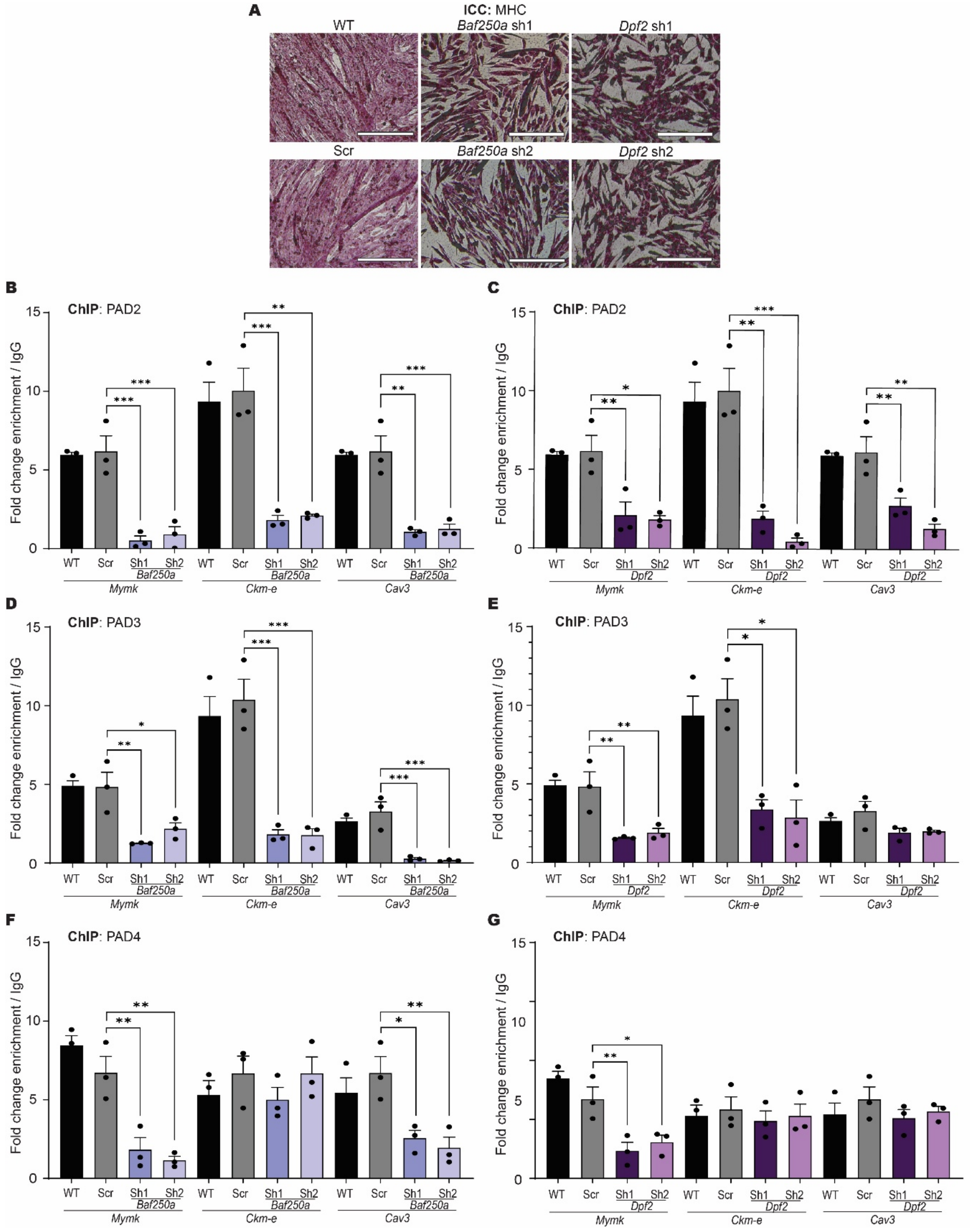

**Supplemental Figure 4: Knockdown of the genes encoding cBAF-specific mSWI/SNF chromatin remodeling enzyme subunits BAF250A and DPF2 reduces PAD enzyme binding to myogenic gene regulatory sequences in differentiating myoblasts. (A)** *Baf250A* or *Dpf2* knockdown in primary myoblasts impaired differentiation as demonstrated by immunostaining cells for MHC 24 h post-differentiation. **(B,C)** ChIP experiments for PAD2 at the indicated myogenic gene regulatory sequences in primary myoblasts differentiated for 24 h following shRNA-mediated knockdown of *Baf250A* or *Dpf2*, respectively. **(D,E)** ChIP experiments for PAD3, as described in (B,C). **(F,G)** ChIP experiments for PAD4, as described in (B,C). Data presented here was generated side-by side with data presented in Fig. 6 but presented separately to simplify Fig. 6. Distinct shRNAs targeting each *Padi* gene are indicated by Sh1 and Sh2. Bar graphs represent the mean ± SEM from three independent experiments. *p < 0.05, **p < 0.01, ***p < 0.001, and ****p < 0.0001. WT, Wild-type; Scr, scrambled sequence shRNA.

### Supplemental figure 5

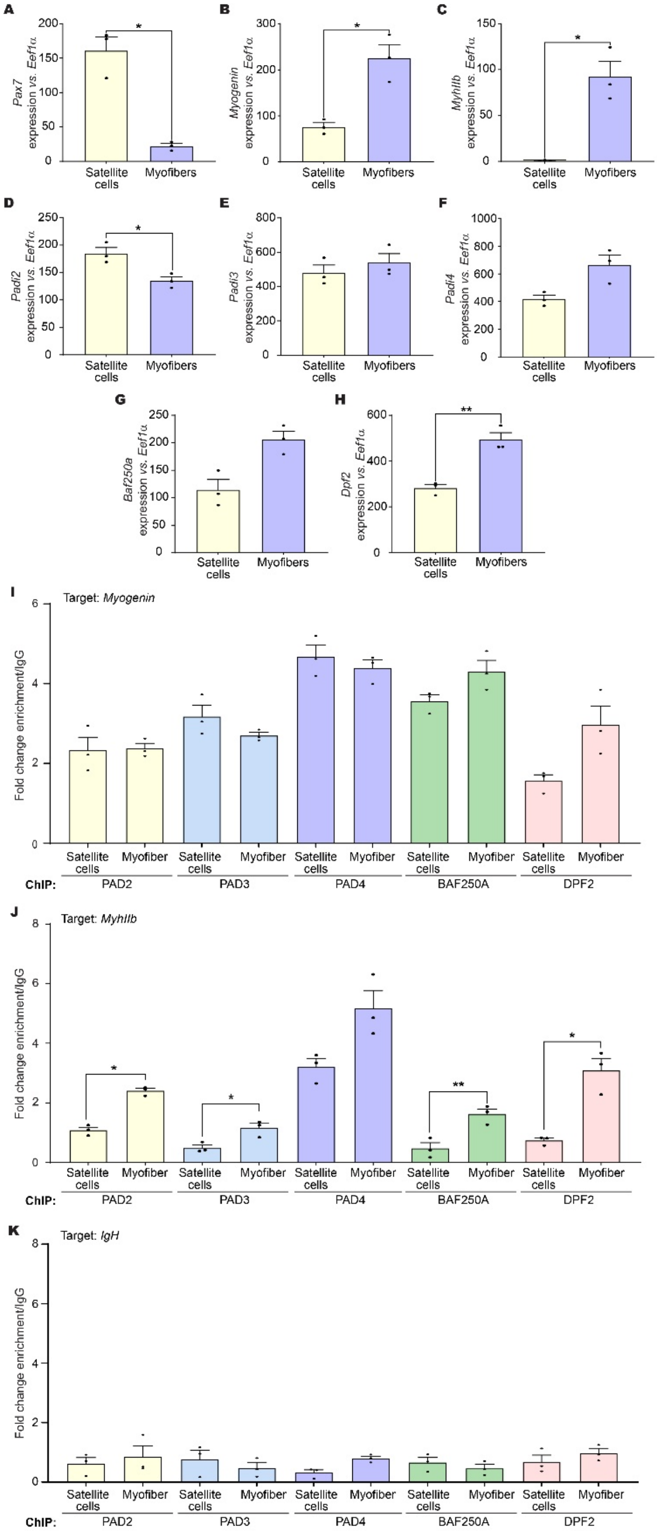


**Supplemental Figure 5: PAD enzyme and cBAF-specific subunits of mSWI/SNF chromatin remodeling enzymes bind to myogenic gene regulatory sequences in**

**murine satellite cells and myofibers. (A-H)** Stable mRNA levels in satellite cells and myofibers isolated from mouse TA muscle were determined by qPCR for *Pax7* (A), *myogenin* (B), *MyhIIb* (C), *Padi2* (C), *Padi3* (D), *Padi4* (E), *Baf250A* (F), and *Dpf2* (G). **(I-K)** ChIP experiments for the indicated proteins at the myogenin promoter (I), MyhIIB promoter (J), and IgH enhancer (K) as a negative sequence control. Bar graphs represent the mean ± SEM from three independent experiments. *$p < 0.05$, **$p < 0.01$, ***$p < 0.001$, and ****$p < 0.0001$.

#### Supplementary figure 6

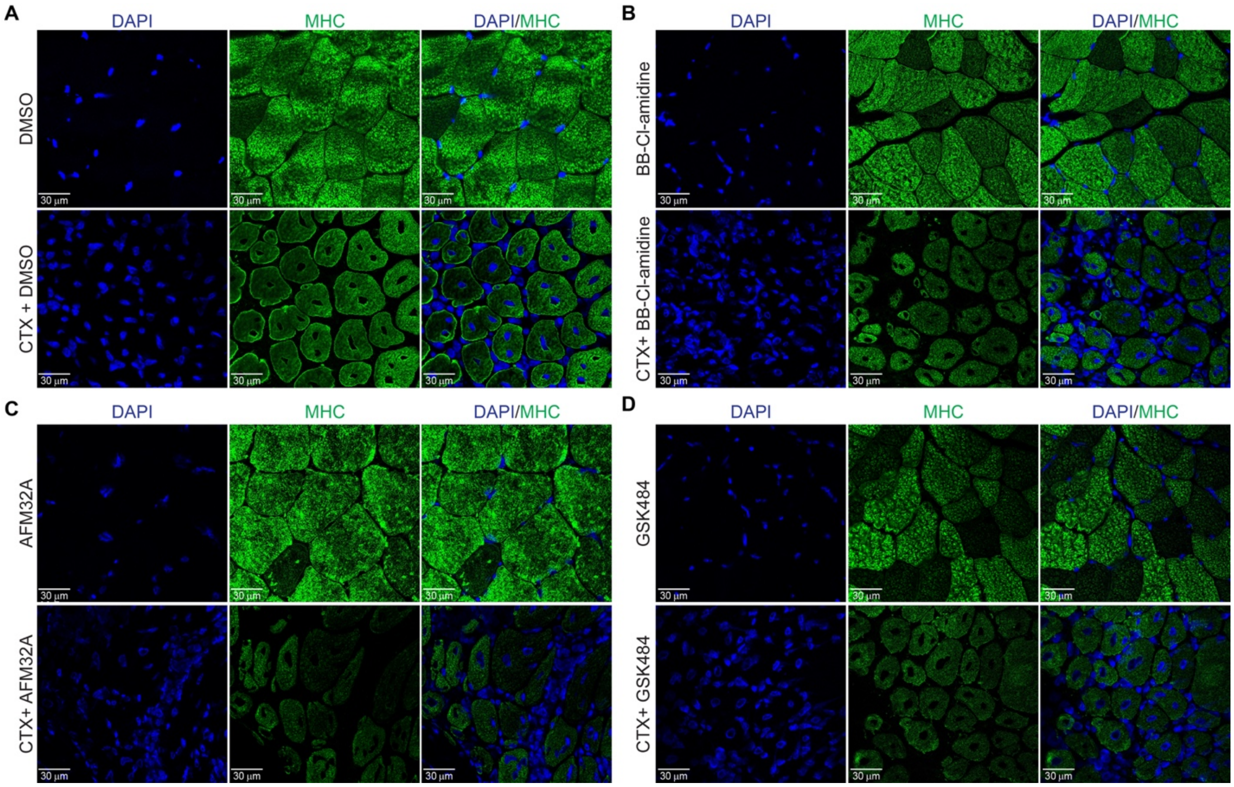


**Supplementary Figure 6: MHC expression is reduced in regenerating TA muscle following injury when the regenerating muscle is treated with PAD inhibitors. (A-D)** Confocal micrographs of MHC immunostaining (green) of TA muscle from uninjured or injured male mice injected with DMSO (A), BB-Cl amidine (B), AFM32a (C), or GSK484 (D). Nuclei were counterstained with DAPI (blue). Scale bar, 30 µm.

**Supplemental table 1.** Sequences of shRNA used in this study.

| Orig. Target Gene | Clone ID | Target Seq |
|---|---|---|
| *Padi2 Sh1* | TRCN0000101440 | GCACAGAATCTTCCAGAAATA |
| *Padi2 Sh2* | TRCN0000101441 | GCGCTCTTCAAGATGGATGAA |
| *Padi3 Sh1* | TRCN0000101535 | GCGCTGTTGATGGTTAGGAAA |
| *Padi3 Sh2* | TRCN0000101536 | CCAGAGCCTTATCAACTTTAA |
| *Padi4 Sh1* | TRCN0000101830 | CAAGCGAGTTATGGGTCCAAA |
| *Padi4 Sh2* | TRCN0000101833 | AGGAGAATATAGATGACCAAT |

**Supplemental table 2.** Sequences of primers used in this study

| Primer name | 5'→3' Sequence | Use |
|---|---|---|
| q*Pax7* F | GCAGCTGGAGGAGCTAGAGAAG | Gene expression |
| q*Pax7* R | GTCTCCTGGCTTGATGGAGTC | Gene expression |
| q*Myogenin* F | CAAGTGTGCACATCTGTTCTAGTCTCT | Gene expression |
| q*Myogenin* R | GTATCATCAGCACAGGAGACCTTGGT | Gene expression |
| q*Ckm* F | CTGTCCGTGGAAGCTCTCAACAGC | Gene expression |
| q*Ckm* R | TTTTGTTGTCGTTGTGCCAGATGCC | Gene expression |
| q *MyHCIIb* F | TCAATGAGATGGAGATCCAGCTGAAC | Gene expression |
| q *MyHCIIb* R | GTCCAGGTGCAGCTGTGTGTCCTTC | Gene expression |
| q*Cav3* F | TCAATGAGGACATTGTGAAGGTAGA | Gene expression |
| q*Cav3* R | CAGTGTAGACAACAGGCGGT | Gene expression |
| q*Dpf2* R | TGCCTGTGACATTTGTGGAA | Gene expression |
| q*Dpf2* F | GCCATCATCACAGGGGTGAA | Gene expression |
| q*Mymk* R | GGCAAAGGTTTCTCCCATGCC | Gene expression |
| q*Mymk* F | GTCGGCCAGTGCCATCAGGGA | Gene expression |
| q*Padi2* R | GGTGAAAGGTCAGCCAAGAAG | Gene expression |
| q*Padi2* F | GCCGCCTATACGGGAAAATATG | Gene expression |
| q*Padi3* R | CCCCTAGCAATGACCTCAAC | Gene expression |
| q*Padi3* F | ATAGGCCAGGGGCAAATG | Gene expression |
| q*Padi4* R | GAGCAAGGATGGCCCAAGG | Gene expression |
| q*Padi4* F | GACAGTTCCACCCCAGTGAT | Gene expression |
| q*EEf1A1α* F | AGCTTCTCTGACTACCCTCCACTT | Gene expression |
| q*EEf1A1α* R | GACCGTTCTTCCACCACTGATT | Gene expression |
| *Pax7 A* ChIP F | GTGGCGACAAGGAAGTTCAAACAAAC | ChIP |
| *Pax7 A* ChIP R | AAAGAAAGCCACTCCGCAACCTCTG | ChIP |
| *Myog* ChIP F | ACACCAACTGCTGGGTGCCA | ChIP |
| *Myog* ChIP R | GAATCACATGTAATCCACTGG | ChIP |
| *MyhIIb* ChIP F | CACCCAAGCCGGGAGAAACAGCC | ChIP |
| *MyhIIb* ChIP R | GAGGAAGGACAGGACAGAGGCACC | ChIP |
| *Cav3* ChIP F | CCTAGGTGTCTCAGTCCAGTTA | ChIP |
| *Cav3* ChIP R | CTGCCACGTAGATCTTGGAAAT | ChIP |
| *Mck-e* ChIP F | GACACCCGAGATGCCTGGTT | ChIP |
| *Mck-e* ChIP R | GATCCACCAGGGACAGGGTT | ChIP |
| *Mymk* ChIP F | GGTGGAGAAGGTCAGCTTGG | ChIP |
| *Mymk* ChIP R | TGGCCATGTCCTTTGTCCTG | ChIP |
